# Heavy-chain CDR3-engineered B cells facilitate *in vivo* evaluation of HIV-1 vaccine candidates

**DOI:** 10.1101/2022.11.07.515497

**Authors:** Wenhui He, Tianling Ou, Nickolas Skamangas, Charles C. Bailey, Naomi Bronkema, Yan Guo, Yiming Yin, Valerie Kobzarenko, Xia Zhang, Andi Pan, Xin Liu, Ava E. Allwardt, Debasis Mitra, Brian Quinlan, Rogier W. Sanders, Hyeryun Choe, Michael Farzan

**Author notes:** Equal authors (Wenhui He, Tianling Ou).

## Abstract

V2-glycan/apex broadly neutralizing antibodies (bnAbs) recognize a closed quaternary epitope of the HIV-1 envelope glycoprotein (Env). This closed structure is necessary to elicit apex antibodies and useful to guide maturation of other bnAb classes. To compare antigens designed to maintain this conformation, apex-specific responses were monitored in mice engrafted with a diverse repertoire of B cells expressing the HCDR3 of the apex bnAb VRC26.25. Engineered B cells affinity matured, guiding improvement of VRC26.25 itself. We found that soluble Env (SOSIP) variants differed significantly in their ability to raise anti-apex responses. A transmembrane SOSIP (SOSIP-TM) delivered as an mRNA-lipid nanoparticle elicited more potent neutralizing responses than multimerized SOSIP proteins. Importantly, SOSIP-TM elicited neutralizing sera from B cells engineered with the predicted VRC26.25-HCDR3 progenitor, which also affinity matured. Our data show that HCDR3-edited B cells facilitate efficient *in vivo* comparisons of Env antigens and highlight the potential of an HCDR3-focused vaccine approach.

## INTRODUCTION

HIV-1 infected persons can raise potent broadly neutralizing antibodies (bnAbs) (Walker et al., 2009; Wu et al., 2010; Liao et al., 2013), but to date no vaccine strategy can do so consistently. These bnAbs nonetheless serve as important guides to the design of HIV-1 immunogens by identifying conserved target epitopes and highlighting features of B-cell receptors (BCRs) necessary to recognize these epitopes (van Gils and Sanders, 2013; Halper-Stromberg et al., 2014; West Jr et al., 2014). Unfortunately, these features often include hard-to-elicit substitutions, insertions, and deletions that emerge only after years of active infection. Thus, sequential immunization strategies are presumed necessary to elicit bnAbs recognizing the CD4-binding site and V3-glycan epitopes of Env (Escolano et al., 2016; Klasse et al., 2016; Steichen et al., 2016; Tian et al., 2016). However, another class of bnAbs, those recognizing the V2-glycan/apex epitope, have qualitatively different properties that may simplify their elicitation (McLellan et al., 2011; Andrabi et al., 2015; Lee et al., 2017; Voss et al., 2017). For example, apex bnAbs require relatively less somatic hypermutation (SHM) and much of their binding energy is localized to their heavy-chain complementarity-determining 3 regions (HCDR3s). These HCDR3s are unusually long, acidic, and tyrosine-sulfated (Andrabi et al., 2015; Lee et al., 2017). Antibodies with similarly long HCDR3s and tyrosine-sulfation motifs are readily observed in the repertoires of HIV-1 negative persons (Briney et al., 2019), and these antibodies might serve as apex bnAb precursors. Consistent with this supposition, apex bnAbs emerge more quickly and frequently after infection than other bnAb classes (Walker et al., 2010; Georgiev et al., 2013; Roark et al., 2021). Moreover, although their breadth is relatively limited, apex bnAbs are typically more potent than other bnAbs (McLellan et al., 2011; Pancera et al., 2013; Doria-Rose et al., 2014; Doria-Rose et al., 2016). Thus, the apex epitope remains a critical target of efforts to develop an effective HIV-1 vaccine.

Apex bnAbs such as PGT145 and VRC26.25 also serve an additional function in vaccine design. Specifically, because they recognize only properly assembled Env trimers in a closed conformation (Sok et al., 2014; Andrabi et al., 2015; Sanders et al., 2015), they help purify (Ringe et al., 2015; Cupo et al., 2019) and quality control candidate Env antigens (Richardson et al., 2018). Env trimers in this closed state hide otherwise immunodominant non-neutralizing epitopes and ensure that key neutralizing epitopes are presented as they are on functional virions. Hence, considerable efforts have been applied to increasing the stability and immunogenicity of soluble, trimeric Env antigens, building on SOSIP (Sanders et al., 2013; Guenaga et al., 2015; Steichen et al., 2016; Guenaga et al., 2017; Medina-Ramírez et al., 2017) and native-flexible linked (NFL) architectures (Georgiev et al., 2015; Sharma et al., 2015; Martinez-Murillo et al., 2017), among others (Aldon et al., 2018; He et al., 2018). Rabbits and macaques immunized with these trimers can elicit autologous and weak heterologous neutralizing responses, sometimes including detectable anti-apex responses (Klasse et al., 2016; McCoy et al., 2016; Voss et al., 2017; Brouwer et al., 2019; Sliepen et al., 2019).

Efforts to design and improve Env immunogens that maintain this closed apex-bnAb-binding conformation nonetheless face challenges. The quaternary apex epitope is difficult to maintain during protein purification, and nearly impossible to monitor *in vivo*. The apex bnAbs PGT145 (Ringe et al., 2015; Cupo et al., 2019) or VRC26.25 (Chuang et al., 2017) are used to enrich for properly assembled trimers, but the structural integrity of these trimers can vary during subsequent handling or after immunization. The emergence of mRNA vaccines can bypass the need for protein *in vitro* handling (Pardi et al., 2020; Saunders et al., 2021; Zhang et al., 2021), but *in vivo* stability remains a key variable in antigen design. However, the stability of these antigens is not the only determinant of their ability to raise potent bnAbs. For example, immunodominant non-neutralizing epitopes can compete with neutralizing ones (Doms and Moore, 2000; Sanders et al., 2013; de Taeye et al., 2015), cellular and serum proteases can destroy key epitopes, non-neutralizing antibodies can drive disassembly of trimeric antigens, and unanticipated interactions with host-cell proteins or extracellular matrix elements can limit their access to follicular dendritic cells (Turner et al., 2021; van Schooten et al., 2021).

Thus, efficient and physiologically relevant *in vivo* systems for measuring anti-apex responses will be critical for developing better Env antigens. However, current wild-type rodent, rabbit, or macaque models are not optimal in large part because the diversity (D) gene segments, key contributors to the HCDR3, are highly species-specific (Lefranc, 2014; Lefranc et al., 2015).Transgenic mice can be engineered to express mature or progenitor apex bnAbs (Crooks et al., 2021; Melzi et al., 2022), but these mice are slow to generate or modify, and their antigen reactive repertoires are essentially monoclonal, biasing antigen comparisons. Strategies to engineer mature murine B cells to express human bnAbs and adoptively transfer these cells into wild-type mice have been developed for novel cell-based therapies (Voss et al., 2017; Hartweger et al., 2019; Moffett et al., 2019; Voss et al., 2019; Huang et al., 2020; Nahmad et al., 2020; Nahmad et al., 2022). These engineered B cells proliferate in response to antigen and generate neutralizing sera, but this response has impaired somatic hypermutation, and is monoclonal and therefore unrepresentative of a human repertoire. Thus, there remains a need for a robust, sensitive, and adaptable system for monitoring the stability and immunogenicity of Env trimers *in vivo*.

Here we demonstrate that a diverse repertoire of murine BCRs engineered to express the HCDR3s of several apex bnAbs can bind soluble Env trimers. When B cells modified to express the VRC26.25 HCDR3 were introduced into wild-type mice, they proliferated, class switched, hypermutated, and generated potent neutralizing sera following immunization with a range of Env immunogens. Notably, these engineered B cells affinity matured, as indicated by the ability of hypermutated HCDR3 to improve the potency of wild-type VRC26.25. Using this system, multiple SOSIP variants were evaluated *in vivo* for their antigenicity. Among them, a version of the chimeric SOSIP protein raised the most potent apex-antibody responses and relatively few non-neutralizing antibodies. Responses to these SOSIP protein variants were markedly enhanced when they were delivered as an mRNA vaccine and expressed as transmembrane proteins (SOSIP-TM). This more immunogenic vaccine candidate induced neutralizing responses from B cells expressing the HCDR3 of the apex bnAbs PG9 and PG16, as well as affinity maturation of the predicted unmutated common ancestor (UCA; (Doria-Rose et al., 2014) of the VRC26.25 HCDR3. Thus, the approach taken here can accelerate development of stable Env immunogens that elicit apex bnAbs from human HCDR3 precursors.

## RESULTS

### Mice engrafted with HCDR3-edited B cells generate neutralizing sera after immunization

We have previously demonstrated that primary human B cells can be edited to express an exogenous HCDR3 as part of an otherwise diverse native human antibody repertoire (Ou et al., 2022). Specifically, we developed techniques to introduce the HCDR3s of the apex bnAbs PG9 and PG16 into naïve mature B-cell receptors encoded by VH-1, VH-3, and VH-4 families of variable genes. Reflecting the high dependence of apex bnAbs on their extended, usually tyrosine-sulfated, HCDR3s (**Fig. 1A**), we showed that B cells engineered to express these HCDR3s bound SOSIP antigens in the context of diverse human heavy- and light chains. These initial studies in human cells suggested that B cells so edited might expand and affinity mature *in vivo* in response to HIV-1 immunogens. To test this hypothesis, primary murine B cells were modified using the same strategy. Specifically, splenic B cells were electroporated with ribonucleoproteins (RNP) composed of the Mb2Cas12a CRISPR effector protein and guide RNA (gRNA) complementary to a region within the JH4 segment (**Fig. 1B**). Mb2Cas12a efficiently recognizes a convenient GTTC PAM present in a usually conserved region of JH4, allowing it to cut a site near the DJ junction. A short single-stranded homology-directed repair template (HDRT), with a 5’ homology arm complementing a ∼70-nucleotide consensus murine VH1 sequence and a 3’ homology arm complementing ∼60-nucleotide 3’-region of JH4 and its adjacent intron, was used to introduce the full HCDR3 of several apex bnAbs, or a hemagglutinin (HA) tag sequence, into the VDJ-recombined murine variable gene. HDRTs were modified with two 3’ phosphorothioates that markedly enhanced editing efficiency (**Fig. S1A-B**). B cells were activated with LPS (Huang et al., 2020), which enhanced editing efficiencies comparably to other reported activation protocols (Hartweger et al., 2019; Nahmad et al., 2020) (**Fig. S1C**). Using this approach, we introduced the HCDR3 regions of the apex bnAbs VRC26.25, PG9, PG16, and the predicted unmutated common ancestor (UCA) of the VRC26-family (VRC26-UCA) into primary mouse B cells and measured their abilities to bind an HA-antibody control or a SOSIP trimer (**Fig. 1C-D**). B cells edited to express the HA-tag as a HCDR3 were used as negative controls. We observed that this SOSIP trimer specifically bound B cells edited to express all four apex-bnAb-derived HCDR3s, and that it bound cells expressing the VRC26.25 HCDR3 most efficiently.

**Figure 1.**
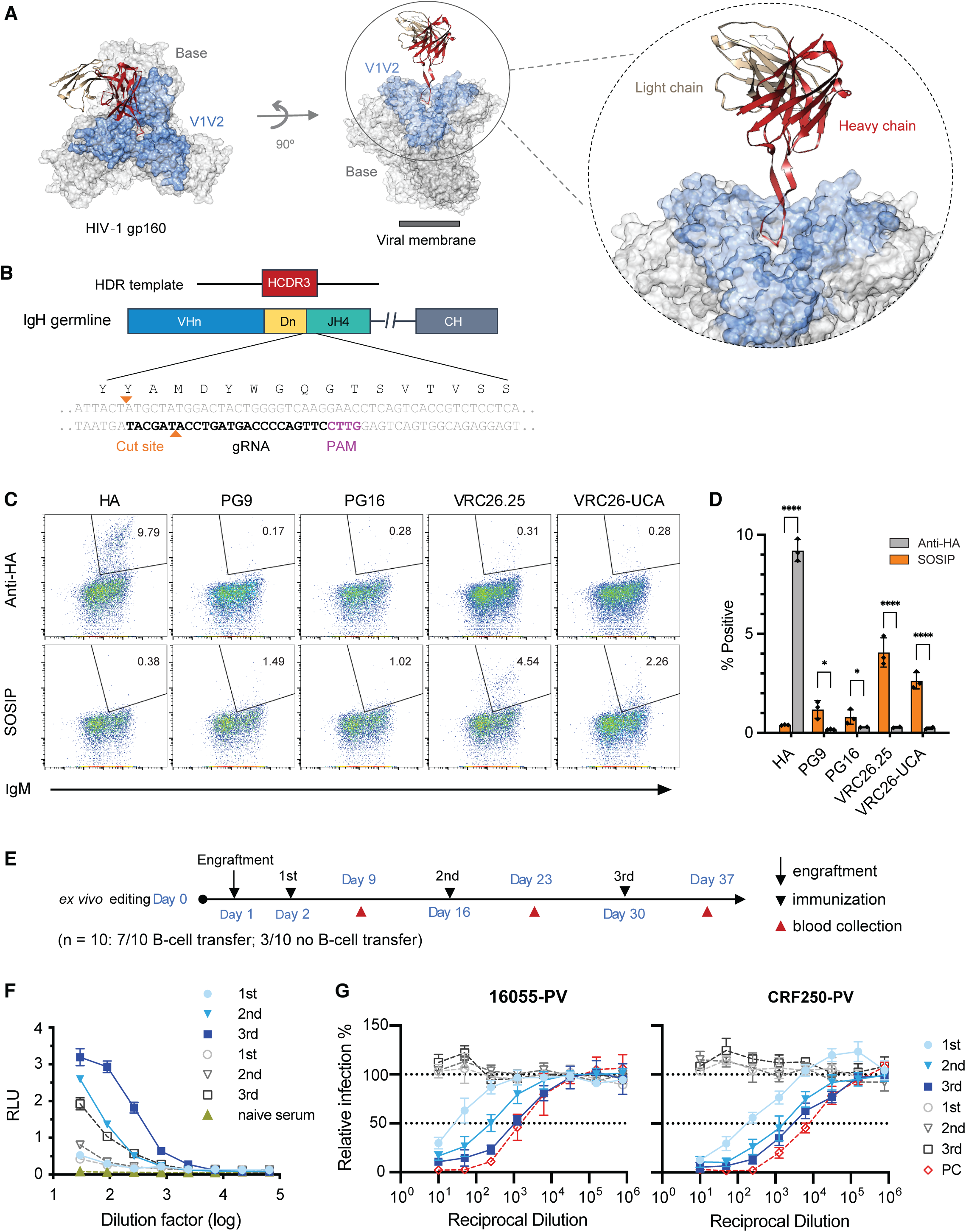
Mice engrafted with HCDR3-edited B cells generate neutralizing sera after immunization. (**A**) A structure of the VRC26.25 Fab bound to an Env trimer (PDB: 6VTT, (Gorman et al., 2020)). The VRC26.25 heavy-chain variable chain is indicated in red, with the light-chain variable chain in tan. Light blue indicates the V1V2 region formed by the V1 and V2 Env variable loops, where the apex epitope locates. Light grey color indicates the base region. Note that the long HCDR3 protrudes into a basic cavity formed by all three Env protomers. (**B**) The coding region of the murine antibody heavy-chain variable (VH) locus is represented. The HCDR3 is encoded in VDJ-recombined heavy chain by the 3’ end of a VH gene (blue), one or more D genes (yellow), and the 5’ end of the JH4 gene (green). The gRNA of the CRISPR Mb2Cas12a effector protein complements the conserved 3’ of JH4, while cleaving near the site of HCDR3 insertion for efficient editing. As indicated, JH4, the most 3’ of murine JH genes, contains optimally positioned non-canonical PAM sequence (GTTC), efficiently cleaved by the Mb2Cas12a ortholog. This PAM, the gRNA, and the Mb2Cas12a cut sites are indicated. At the top of the figure, the homology-directed repair template (HDRT) encoding an exogenous HCDR3 (red) is shown, bounded by two short homology arms (60 to 72 nucleotides). The 5’ homology arm complements a VH1 consensus sequence and the 3’ arm complements a conserved region of JH4 and adjacent intronic sequences. (**C**) The HCDR3s of the apex bnAbs PG9, PG16, VRC26.25, VRC26-UCA, or a hemagglutinin (HA) tag were introduced into the BCRs of primary murine B cells and analyzed by flow cytometry for their ability to bind a fluorescently an anti-HA antibody or fluorescently labeled SOSIP trimer whose apex is formed by the clade B 16055 isolate (vertical axis). The horizontal axis indicates expression of surface IgM, and its loss indicates imprecise non-homologous end joining (NHEJ) after Mb2Cas12a-mediated cleavage. Number within each panel indicates the percentage of cells that bind an anti-HA antibody or SOSIP trimer. (**D**) Results of three independent experiments similar to those shown in panel C are summarized. Error bars indicate SEM. Statistical differences compared to negative controls were determined by two-way ANOVA (*p<0.05; ****p<0.0001). (**E**) A timeline of the B-cell engraftment and immunization protocol used in subsequent panels. B cells isolated from CD45.1-positive mice were engineered to express the VRC26.25 HCDR3 and engrafted at day 1. Mice were immunized with 16055-ConM-v8.1ds SOSIP conjugated to a mi3 60-mer scaffold on days 2, 16, and 30, and blood was harvested at day 9, 23, and 37 for ELISA and neutralization studies. Additional control mice were not engrafted with exogenous B cells but otherwise immunized identically. (**F**) Sera pooled from seven mice, harvested after each indicated immunization (filled blue symbols), were evaluated for their ability to bind 16055-ConM-v8.1ds SOSIP by ELISA. Three unengrafted, immunized mice (open grey symbols) served as controls. Naïve sera (filled olive symbol) were obtained from mice without engraftment or immunization. (**G**) The ability of pooled sera from panel **F** to neutralize pseudoviruses bearing the Envs of indicated HIV-1 isolates (CRF250, 16055) is shown. Naïve sera mixed with wild-type VRC26.25 antibody to 2 µg/ml is used as a positive control.

To determine whether these cells could respond to antigen *in vivo*, we isolated B cells from the spleens of B6 CD45.1 mice, edited them as above to introduce the VRC26.25 HCDR3 region, and adoptively transferred them to wild-type (CD45.2) C57BL/6J mice. Recipient mice initially received 30 million cells and then were immunized with adjuvanted SOSIP trimers conjugated to a mi3 60-mer scaffold (Bruun et al., 2018; Keeble et al., 2019) according to the schedule represented in **Fig. 1E**. Serum harvested one week after each of three immunizations was characterized for its ability to bind SOSIP trimers (**Fig. 1F**) and to neutralize pseudoviruses presenting the Envs of the CRF250 and 16055 isolates (**Fig. 1G**). We observed that mice produced increasing SOSIP-binding antibodies with each immunization. The neutralization efficiency increased similarly in mice engrafted with edited B cells. In contrast, mice that did not receive edited cells, but which were immunized on the same schedule, did not raise neutralizing sera, indicating that that edited B cells were responsible for the neutralizing activity.

### HCDR3-edited B cells migrate to germinal centers, class switch, and hypermutate

We next sought to determine if edited B cells migrated to germinal centers in the lymph nodes and spleens of recipient mice. An experiment similar to that shown in **Fig. 1** was performed (**Fig. 2A**) except that spleens and lymph nodes were harvested from two mice after each of three immunizations, and control mice were also engrafted with cells expressing the VRC26.25 HCDR3, but not immunized. We observed an increase in germinal center (CD38-/GL-7+) B cells, including both donor CD45.1+ cells and recipient CD45.1-cells, compared to non-immunized mice (**Fig. 2B and C**). The percentage of SOSIP binding donor cells [SOSIP(+)/CD45.1+] increased after each immunization, whereas the CD45.1+/SOSIP(-) cells did not (**Fig 2B and D**), suggesting that only antigen-reactive cells proliferated in germinal centers.

**Figure 2.**
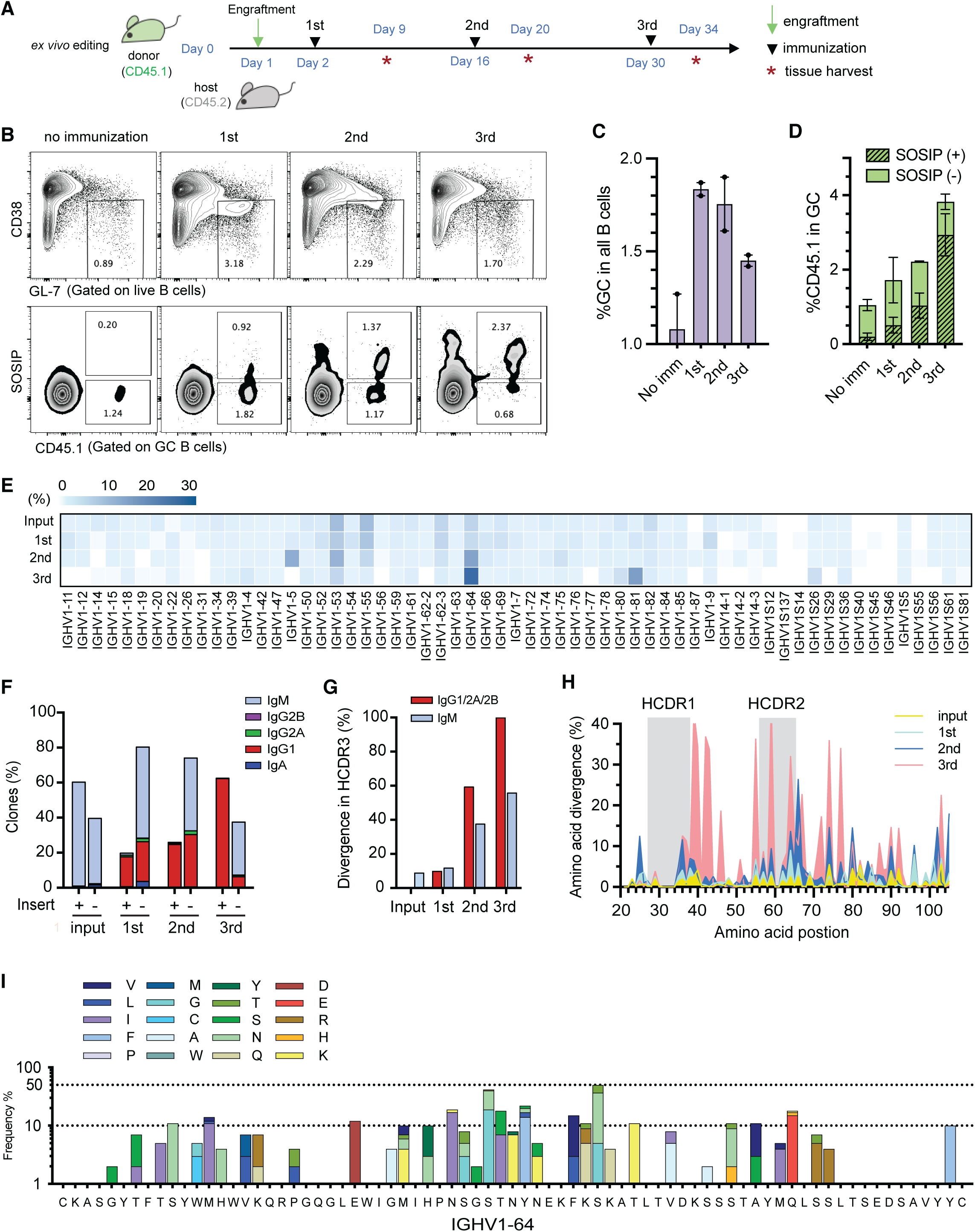
HCDR3-edited B cells migrate to germinal centers, class switch, and hypermutate. (**A**) A timeline of studies used in this figure. Host mice distinguished by a CD45.2 marker were adoptively transferred at day 1 with VRC26.25 HCDR3-edited B cells derived from a CD45.1 donor. Six engrafted mice were immunized with SOSIP (BG505-ConM-v8.1ds) antigens conjugated to a mi3-60mer scaffold at days 2, 16, and 30. Spleens and lymph nodes were harvested from two mice each at the indicated time points (red asterisk) and SOSIP-bound donor B cells were sorted for next-generation sequencing. (**B**) B cells isolated from spleen and lymph nodes were pooled after the indicated immunization, or from engrafted mice without immunization, were analyzed by flow cytometry. A representative flow figure is shown for each group. Germinal center (GC) B-cells identified as GL7+/CD38-were gated and quantified (top panels). These gated cells were then analyzed for the frequency of CD45.1+, indicating they derived from donor mice, and for SOSIP binding. Numbers in bottom panels indicate percentages of SOSIP(+) and SOSIP(-) donors B cells found in GC, respectively. (**C**) The mean percentage of GC cells from two mice harvested after each indicated immunization is shown. (**D**) The percentage of CD45.1 donor cells gated from GC is shown, with SOSIP(+) (crosshatches) and SOSIP(-) cells (solid) indicated. Each dot in (**C**) and (**D**) represents percentage analyzed from one mouse. (**E**) The VH1 gene segments of donor CD45.1 cells isolated from mice described in panels **A-D** were analyzed by next-generation sequencing (NGS). SOSIP-binding donor B cells [(CD45.1+/ SOSIP(+)] were isolated from two mice at each time point and pooled for sequencing. The diversity and frequency of VH1 gene segments of successfully edited donor cells is represented with a heat map as percentage of total clones for the indicated time points. ‘Input’ indicates CD45.1+/SOSIP(+) cells after editing but before engraftment. Note the persistence of repertoire diversity but with enrichment for specific variable chains. (**F**) The distribution of isotypes of CD45.1+ donor cells with (insert +) or without (insert -) the VRC26.25-HCDR3 is shown. Cells were sorted before engraftment (input) or after each of the indicated immunizations. Note that a higher proportion of cells underwent class-switch recombination in the cells bearing the VRC26.25-HCDR3 insert. (**G**) The percentage of inserted HCDR3 with amino acid changes is shown for IgM and class-switched (IgG1/2A/2B) BCR. (**H**) The frequency of IgH (position 21-105) amino acid mutations in clones expressing the inserted HCDR3 is shown for each immunization. Grey boxes indicate HCDR1 and HCDR2 regions. (**I**) The frequency of VH1-64 (position 21-105) amino acid mutations in clones expressing the inserted HCDR3 is shown after the third immunization. Each amino acid represented with the indicated color. The germline sequence of VH1-64 is shown below the figure.

Using next generation sequencing (NGS), we also investigated the heavy chain diversity of VRC26.25 HCDR3-edited B cells in lymph nodes and spleens of each mouse group (**Fig. 2E**). We observed that this HCDR3 was present in murine BCRs encoded by many VH1-family segments, and that the original distribution of successfully edited VH1 genes was largely reflected in immunized mice. We further observed an enrichment of cells with BCRs from VH1-53, −64, and −81 families over successive immunizations. Similar enrichment was also observed in unedited donor B cells (**Fig. S2A**), suggesting that this enrichment is independent of apex recognition. We conclude that the repertoire diversity of HCDR3-edited B cells is largely preserved *in vivo* after multiple immunizations, consistent with the predominant contribution of the VRC26.25 HCDR3 to Env association.

We also investigated class-switching and hypermutation frequencies of HCDR3-engineered BCRs. We observed that the number of class-switched CD45.1 donor cells expressing the VRC26.25 HCDR3, but not those without this insert, increased with immunization, and that the majority of these edited cells switched to the IgG1 isotype (**Fig. 2F**). The number of HCDR3s with one or more amino acid mutations also increased with immunization in both IgM and IgG, with more mutations associated with IgG isotypes (**Fig. 2G**). The frequency of framework, HCDR1, and HCDR2 mutations also increased with immunization in HCDR3-edited donor cells (**Fig. 2H**), with diverse amino acid changes emerging at multiple sites (**Fig. 2I**). Collectively, these data show that HCDR3-edited B cells largely maintain their underlying diversity while they proliferate, class switch, and hypermutate in response to immunization.

### Affinity maturation of HCDR3-edited B-cell receptors

We analyzed mutation patterns in the engineered HCDR3 regions after each immunization (**Fig. 3A**), and again observed repeated amino acid changes at multiple locations, and most notably a Q100nE mutation. It was observed in other VRC26 variants from the human donor CAP256 and was previously shown to improve the breadth and potency of VRC26.25 (Doria-Rose et al., 2014; Yin et al., 2021). This mutation emerged in all six mice we characterized, including those immunized only once, and the frequency was increased by 25- or 45-fold after receiving two or three immunizations, respectively. This observation was also repeated in six additional mice in a parallel experiment using a keyhole limpet hemocyanin (KLH) carrier protein to present SOSIP proteins (**Fig. S2B**), which are less immunogenic due to the inconsistent conjugate orientations. We analyzed individual HCDR3 mutations that emerged in all three groups of mice (repeated mutations indicated with triangles in **Fig. 3A**) and observed that five of nine characterized mutations increased binding of VRC26.25 to SOSIP proteins generated from two HIV-1 isolates (**Fig. S2C**). Combinations of these mutations further increased their binding, with highest SOSIP-binding observed with triple (NER) and quadruple (NERE) changes (**Fig. S2D**). These increases in binding affinity correlated with neutralization potency against isolates that are difficult to neutralize by VRC26.25 (**Fig. 3B**). We assayed these VRC26.25 variants against a global panel of 13 pseudoviruses. Variants bearing the NER and NERE mutations neutralized these viruses with a geometric mean IC_50_ more than 6-fold lower than the IC_50_ of this already very potent antibody (**Fig. 3C**). We conclude that HCDR3-edited cells affinity mature in response to SOSIP immunization. We further note that mutations that emerge in multiple mice, and the frequency of those increases with immunization, are likely to enhance neutralization potency.

**Figure 3.**
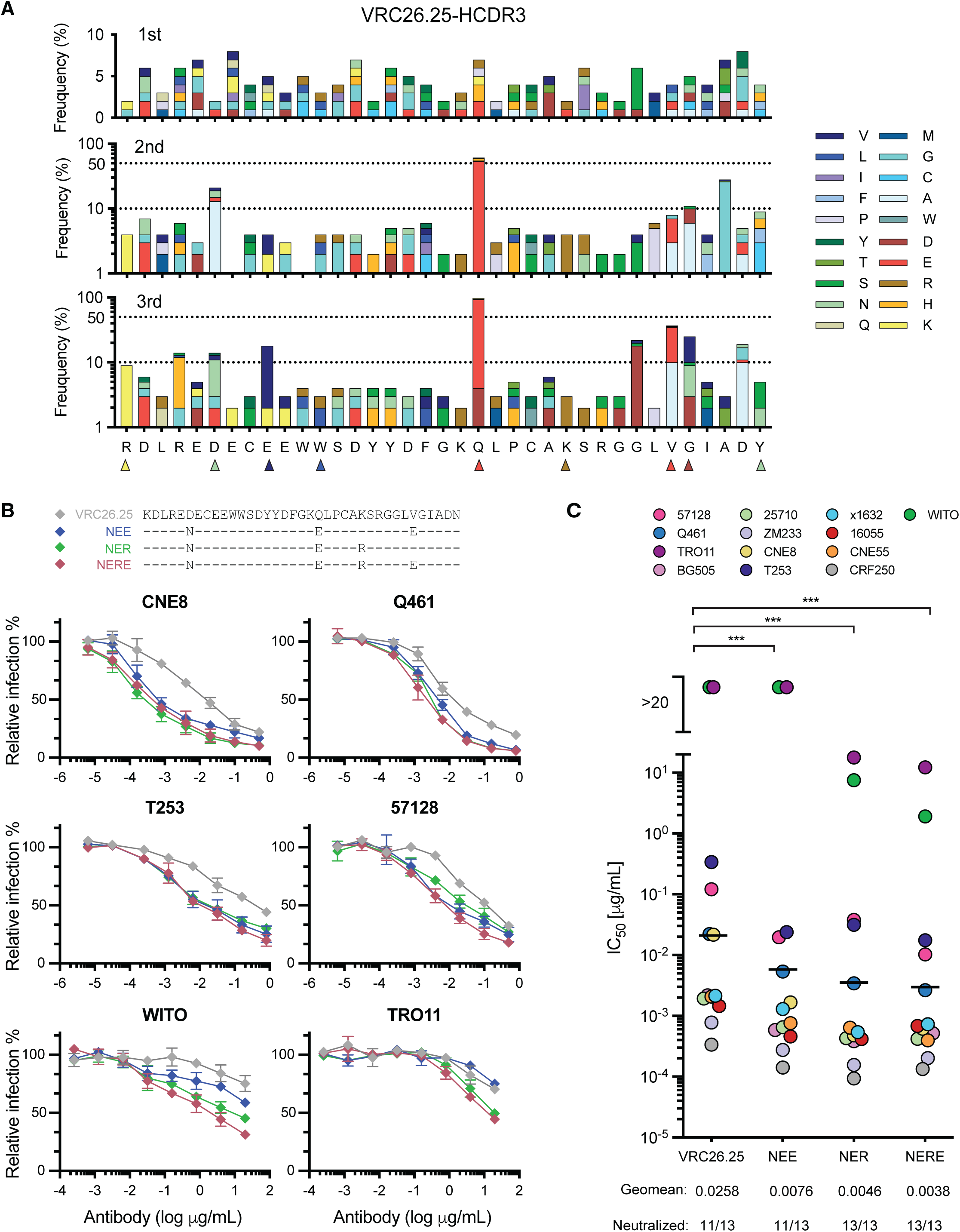
Affinity maturation of HCDR3-edited B-cell receptors. (**A**) The frequencies of amino acid mutations within the HCDR3 region present in HCDR3-edited clones were analyzed after the indicated immunization. The most frequent 9 mutations observed in more than one mouse group represented here, are marked with triangles, with colors indicating the mutated amino acid. These mutations are characterized in subsequent panels. (**B**) Top panel: Combinations of HCDR3 mutations identified in **Fig. S2C** as high affinity binders to ConM-SOSIP (v8.1ds) immunogens bearing either the CRF250 or BG505 V1V2 regions were introduced into full-length, unmodified VRC26.25. Bottom panels: These VRC26.25 variants were characterized for their ability to neutralize the indicated pseudoviruses, including two VRC26.25-resistant isolates (WITO and TRO11). (**C**) The same VRC26.25 variants were used to measure neutralization efficiency of the indicated global panel of HIV-1 isolates. IC_50_ values are represented in colored circles, with geometric mean values indicated by a line. Significant differences from wild-type VRC26.25 were determined by paired t-test (***p<0.001).

### HCDR3-edited B cells facilitate evaluation of SOSIP antigens

We then evaluated whether HCDR3-edited B cells could be used to evaluate candidate Env antigens *in vivo*. A number of SOSIP variants, differing in source isolate, producing cell line, and SOSIP version, were generated (Sliepen et al., 2019; Brouwer et al., 2021a).These constructs are named according to the source isolate of the V1V2 region (BG505, CRF250, ConM, 16055), the source isolate of the SOSIP base (BG505, CRF250, or ConM), the SOSIP version (v5, v7, v8.1) (Sanders et al., 2015; Brouwer et al., 2019; Sliepen et al., 2019; Antanasijevic et al., 2020), and whether an additional disulfide loop (I201C-A433C; “ds”) (Joyce et al., 2017), or additional V3-loop mutations, “mut3” (Chuang et al., 2020), are present. Note that SOSIP versions indicate different stabilizing mutations (**Fig. S3A**) in the base, which is defined as the entire SOSIP trimer subtracting the V1V2 region shown in **Fig. S3B**. We observed that SOSIP proteins with v8.1 mutations, and those produced in N-acetylglucosaminyltranferase I (GnTI)-negative cells and thus lacking complex glycans (Eggink et al., 2010; Sanders et al., 2013), more efficiently bound VRC26.25-HCDR3-edited primary murine B cells (**Fig. S3C-D**).

To compare these antigens *in vivo*, mice engrafted with VRC26.25 HCDR3-edited B cells were immunized up to three times and their sera were harvested one week after each immunization. We first compared three ways to present the ConM-v8.1-ds SOSIP antigen. Specifically, this SOSIP was either conjugated to a KLH carrier, covalently linked via a SpyTag to the Spycatcher-mi3 60-mer (Bruun et al., 2018; Keeble et al., 2019), or introduced as a free SOSIP trimer (**Fig. S4A-B**), each in an MPLA/QuilA mixed adjuvant. We observed significantly higher neutralizing responses against CRF250 pseudoviruses with the mi3-conjugated SOSIP compared to free SOSIP trimer or KLH-conjugated SOSIP (**Fig. S4C**). Thus, all further study of protein antigens used mi3-conjugated SOSIP variants. We also compared 16055-ConM-v8.1ds and BG505-ConM-v8.1ds SOSIP trimers produced in Expi239F cells or in GnTI-negative Expi293F cells. We observed that this SOSIP produced in GnTI-negative cells bound primary HCDR3-edited cells more efficiently (**Fig. S3C-D**) but elicited significantly less potent neutralizing sera (**Fig. S4D**). We compared a series of other conditions and observed that neutralizing responses were relatively independent of the amount of adjuvant or antigen (**Fig. S4E**). Similarly, the method of *ex vivo* activation (LPS or anti-CD180), or the number of adoptively transferred donor cells did not significantly affect neutralization (**Fig. S4F**). Accordingly, in subsequent experiments, LPS was used to activate B cells *ex vivo*, and 25 μg SOSIP-mi3 multimers adjuvanted with 20 μg MPLA was used for immunizations. In most cases, around 10 million cells, were adoptively transferred to a mouse. Since 3-4% of these transferred cells express the VRC26.25 HCDR3 (**Fig. 1C; Fig. S1C**), engineered cells initially comprise less than 0.4% of the host mouse B cells.

We then generated a number of SOSIP variants, each bearing the V1V2 region of the 16055 isolate with a BG505 or ConM base, and directly compared their ability to bind VRC26.25 HCDR3-edited primary murine cells (**Fig. 4A-B**). We focused on the ConM base because it is designed to broaden the antibody responses with the absence of immunodominant holes in the glycan shield (McCoy et al., 2016) and rare, isolate-specific residues (Sliepen et al., 2019). We observed that constructs based on a previously reported v8.1 SOSIP platform bound edited B cells slightly stronger than those based on earlier designs. Further, mice adoptively transferred with edited cells and immunized with v8.1 SOSIPs raised more potent neutralizing responses against all three isolates tested (**Fig. 4C)**. Again, sera from mice that did not receive edited cells did not neutralize any isolate (gray dots), even though anti-SOSIP antibodies could be detected by ELISA in these sera (**Fig. 4D**). Notably, one SOSIP, 16055-ConM-v8.1ds, elicited significantly more binding antibodies in VRC26.25-HCDR3 engrafted mice than in unengrafted mice vaccinated in the same manner, suggesting that this SOSIP raised fewer non-neutralizing murine antibodies (compare grey to colored bars). Differences in neutralization potency correlated with the ability of these sera to compete with VRC26.25 or its inferred germline precursor for binding to the 16055-ConM-v8.1ds SOSIP (**Fig. 4E**). We conclude that SOSIP variants differ in their ability to bind VRC26.25 HCDR3-edited B cells even when their V1V2 regions are identical, and that these variants also differ in their ability to focus immune responses to the V1V2 region. We further observe that SOSIP proteins based on the v8.1ds platform with a ConM base and a 16055 V1V2 region elicited potent apex-focused neutralizing responses. Sera elicited by this SOSIP variant included heterologous neutralizing responses that broadened with each immunization and persisted for more than 200 days (**Fig. S4G-H**). We also compared the ability of different V1V2 regions to elicit apex-antibodies. V1V2 regions of four VRC26.25-sensitive isolates were compared on a ConM-v8.1ds base. Each bound HCDR3-edited B cells with similar efficiencies (**Fig. 4F-G**), but the SOSIP protein expressing the V1V2 region of BG505 generated less potent neutralizing against three pseudoviruses (**Fig. 4H**). Interestingly, these differences were less pronounced than those associated with different SOSIP bases, suggesting that the frequency with which a SOSIP populates a closed conformation is more important than the sequence and structure of these four V1V2 domains. Collectively, these data suggest that HCDR3-edited B cells can provide key insight into the *in vivo* properties of Env antigens.

**Figure 4.**
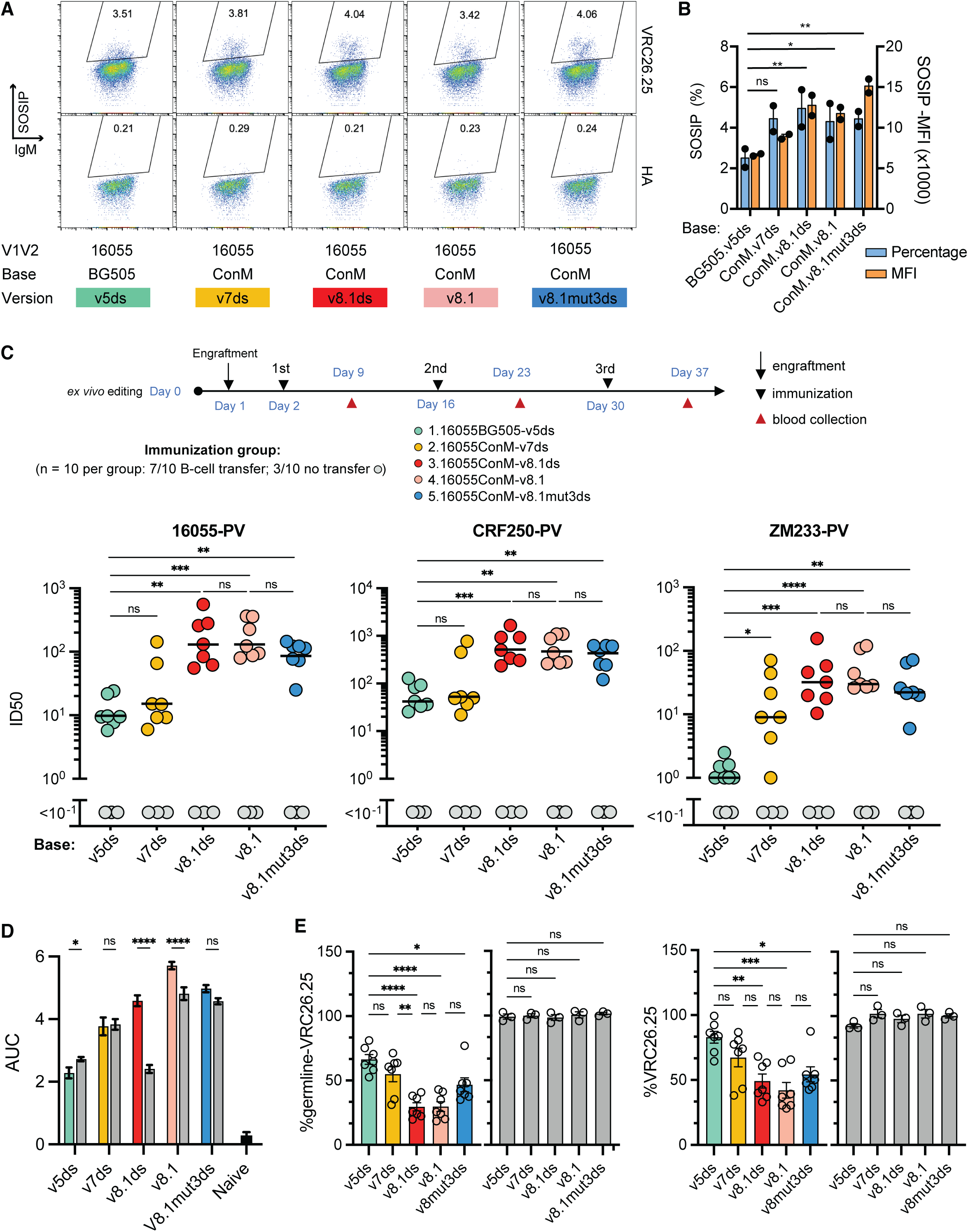
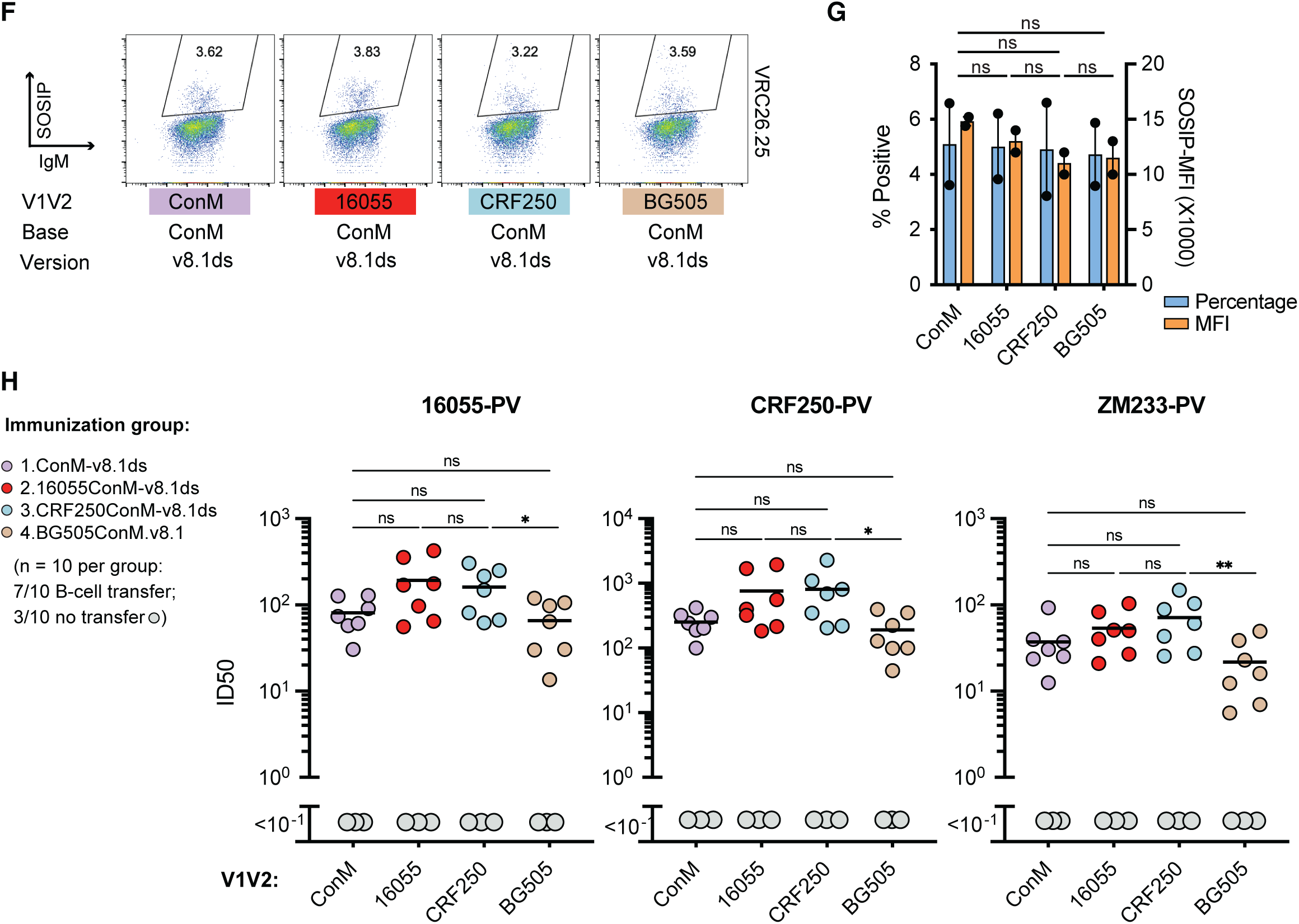
HCDR3-edited B cells facilitate evaluation of SOSIP antigens. (**A**) An example of flow cytometry analysis comparing the binding efficiencies of the indicated SOSIP variants to a population of murine cells edited to express the VRC26.25 HCDR3. Note that all SOSIP proteins included the V1V2 region of the isolate 16055, which replaces the native V1V2 region of the BG505 or ConM SOSIP proteins. Base version indicates sets of previously described mutations designed to stabilize the indicated SOSIP protein (see Fig. **S3A**). SOSIP binding and IgM expression are indicated on the vertical and horizontal axes, respectively, and the percentage of SOSIP-binding cells is indicated within the gate. (**B**) A summary of results from experiments similar to those shown in panel A, with each dot representing an independent experiment. (**C**) Groups of seven mice adoptively transferred with VRC26.25 HCDR3-edited B cells were immunized with the indicated mi3-multimerized SOSIP proteins, as shown at the schedule at the top of the panel. The neutralizing response against the indicated pseudoviruses was measured using serum from each mouse obtained seven days after the second immunization. Lines indicate median 50% inhibitory dilution (ID50), and grey dots indicate results from each of three unengrafted mice immunized in the same manner. (**D**) The ability of sera from these mice to bind to 16055-ConM-v8.1ds SOSIP was determined by ELISA. Colored bars indicate pooled sera from 7 immunized mice engrafted with HCDR3-edited B cells, and gray bars indicated pooled sera from 3 mice immunized without engraftment of edited B cells. Figure shows one of two independent experiments, each with three technical replicates, with similar results. (**E**) Blocking ELISA assays were performed to evaluate the specificity of apex antibody responses. Specifically, the indicated sera analyzed in panel **C** was used to prevent binding of VRC26.25 or the germline form of VRC26.25 to 16055-ConM SOSIP. The percentage of blocking was calculated based on the binding of germline or mature VRC26.25 in the presence of naive mouse serum. Grey indicates sera from immunized mice without engraftment with edited B cells. Each point represents serum from one mouse. (**F-H**) Experiments similar to those shown in panels A-C, except that ConM 8.1 SOSIP variants were modified with V1/V2 regions from the indicated HIV-1 isolates. Significant differences in panels **B-E** and **G-H** were determined by one-way ANOVA (ns, not significant; *p<0.05; **p<0.01; ***p<0.001; ****p<0.0001).

### SOSIP-TM proteins expressed from mRNA vaccines raise more potent neutralizing responses than multimeric protein antigens

Using this mouse model, we compared the immunogenicity of adjuvanted ConM-based SOSIP mi3-multimers to an mRNA-LNP-delivered immunogen expressing the same SOSIP extended through Env residue 712 (HXB2 numbering), thus including the ConM transmembrane (TM) domain and a truncated cytoplasmic region (SOSIP-TM, **Fig. S5A**). Again, VRC26.25 HCDR3-edited B cells were engrafted into recipient mice, which were then immunized with protein SOSIP mi3-multimers or mRNA-LNP expressed SOSIP-TM. In both cases, ConM-v8.1ds SOSIP/SOSIP-TM antigens bearing either the CRF250 or 16055 V1V2 regions were evaluated (**Fig. 5A**). Sera collected 7 days after the second immunization were characterized for neutralization activity against 16055 pseudoviruses. We observed that a 1 μg dose of SOSIP-TM delivered as mRNA-LNP raised comparable neutralizing responses to 25 μg of SOSIP mi3-multimers, whereas 5 μg of SOSIP-TM mRNA-LNP elicited a significantly more potent neutralizing response. Note that 25 μg of adjuvanted protein antigen is a high dose for murine studies (Brouwer et al., 2021b; Gristick et al., 2022), whereas 5 μg is lower than previous studies of mRNA-expressed viral antigens (Laczkó et al., 2020; Melzi et al., 2022; Willis et al., 2022). The most efficient neutralizing response was observed when mice were boosted at 3-week intervals (**Fig. 5A**), and B cells from these mice were further analyzed. We observed that mRNA-LNP immunization induced strong germinal-center reactions (**Fig. S5B**) and robust SHM (**Fig. 5B**). The pattern and rate of SHM was broadly similar to protein immunization with SOSIP mi3-multimers observed in Figure 3A (**Fig. 5B**). We further generated mRNA-LNP expressing four SOSIP-TM variants corresponding to the SOSIP variants characterized in **Fig. 4C**. LNP were quality controlled through dynamic light scattering (**Fig. S5C**) and through cell-surface expression of SOSIP-TM on 293T cells, as measured by the antibodies 2G12 and VRC26.25 (**Fig. 5C-D**). SOSIP-TM variants expressed comparably as indicated by 2G12 binding but varied in their relative binding to VRC26.25, with v8.1 and v8.1ds binding this apex bnAb most efficiently. These same mRNA-LNP were then compared for their ability to raise anti-apex responses *in vivo*. As with proteins antigens, v8.1 and v8.1ds SOSIP-TM variants most efficiently raised apex-specific neutralizing responses (**Fig. 5E**). Sera from the unengrafted mice bound SOSIP trimers comparably to engrafted mice as indicated by ELISA (**Fig. S5D**), but only engrafted mice raised anti-apex responses and these responses correlated with the neutralization activities of these sera (**Fig. S5E**). These data indicate that the advantages of ConM-v8.1 and ConM-v8.1ds persist across platforms and modes of presentation.

**Figure 5.**
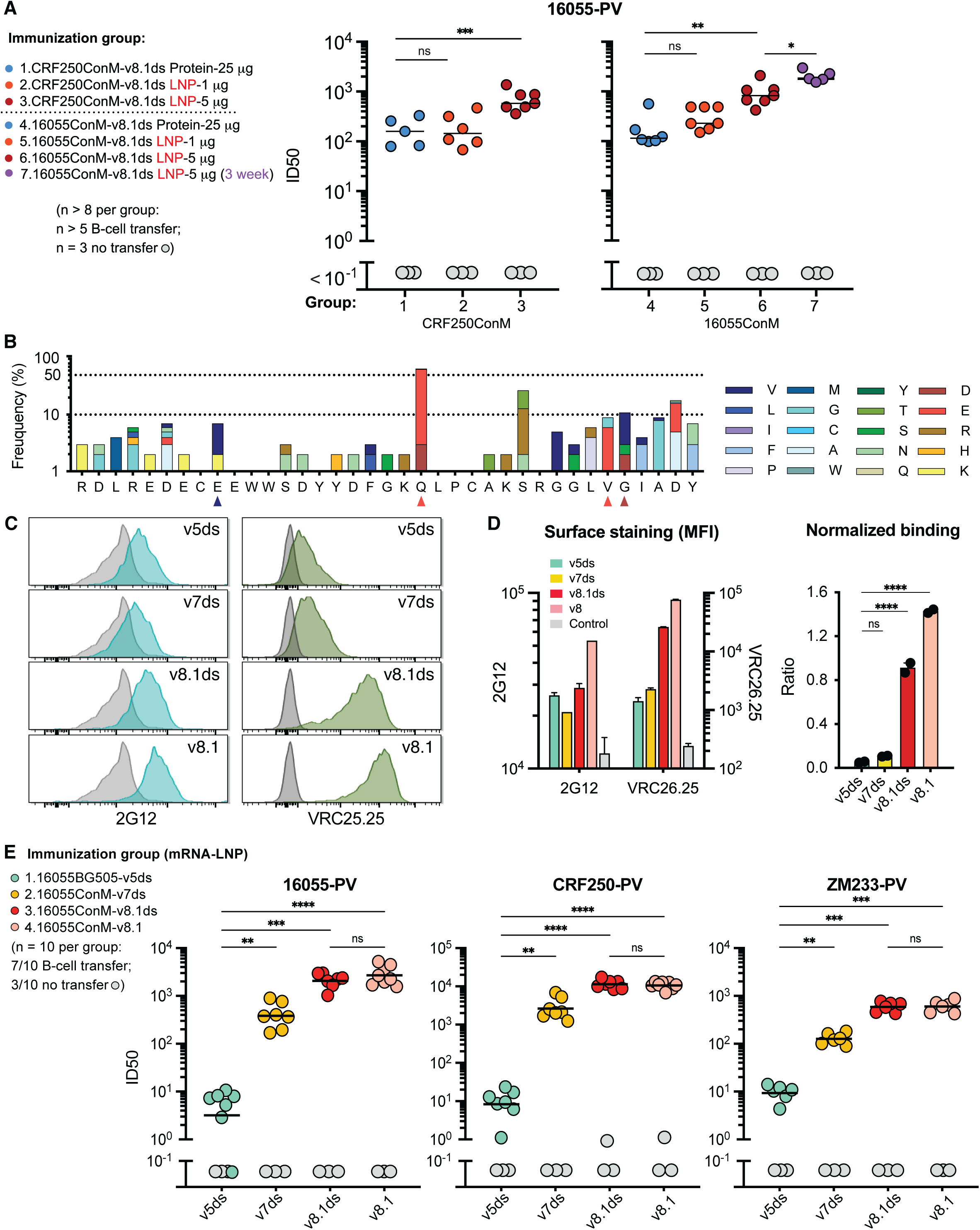
SOSIP-TM proteins expressed from mRNA vaccines raise more potent neutralizing responses than multimeric protein antigens. **(A)** An experiment similar to those performed in Fig. 4C except that the indicated amounts of mRNA-LNP expressing the indicated SOSIP-TM proteins were compared to the soluble forms of the same SOSIP molecules conjugated to the mi3 60-mer scaffold, with prime and boost separated by two or three weeks, as indicated. Sera harvested one week after the boost immunization were characterized. Grey indicates sera from unengrafted mice immunized in the same manner. Statistical difference was determined by one-way ANOVA. **(B)** The frequency of HCDR3 amino acid mutation of donor B cells from immunized mice is shown. Mice were immunized three times with 5 μg mRNA expressing the 16055-ConM-v8.1ds, with each immunization separated by three weeks. **(C)** mRNA-LNP expressing the indicated SOSIP-TM variant were used to transfect 293T cells. Cells were then analyzed by flow cytometry for their ability to bind 2G12, an antibody recognizing conserved gp120 glycans and used here to monitor the overall cell-surface expression, and by VRC26.25. **(D)** A summary of mean fluorescent intensities for experiments similar those shown in panel C is presented at the left, and the ratio of VRC26.25 normalized to 2G12 binding is shown at the right. Panels **C** and **D** are representative of three independent experiments with two technical replicates for each sample. **(E)** An experiment similar to those shown in Figures **4C** and **5A** except that sera from mice vaccinated with mRNA-LNP expressing the indicted SOSIP-TM proteins were compared using the indicated pseudoviruses. Significant differences were determined by one-way ANOVA (ns, not significant; *p<0.05; **p<0.01; ***p<0.001; ****p<0.0001).

### mRNA-LNP elicit neutralizing sera in mice engrafted with B-cells engineered to express the HCDR3s of PG9, PG16, or the VRC26-UCA

All previous experiments used the mature VRC26.25 HCDR3. To determine if a wider range of apex-targeting HCDR3s could also respond to SOSIP-TM immunization, we engrafted B cells with the HCDR3s of the apex bnAbs PG9, PG16, and CH01, as well as the VRC26-UCA. Mice were immunized with mRNA-LNP encoding SOSIP-TM variants of 16055-ConM-v8.1 or CRF250-ConM-v8.1ds. Neutralization responses against the CRF250 or 16055 pseudoviruses were observed after three immunizations in mice engrafted with the PG9, PG16, and VRC26-UCA HCDR3 (**Fig. 6A**). No responses were observed using the HCDR3 of CH01, consistent with its lower affinity and greater dependence on non-HCDR3 elements. Notably, PG9, PG16, VRC26.25, and the VRC26-UCA, but not CH01, share a common tyrosine-sulfated YYDF motif derived from the D3-3 gene segment (**Fig. S6A**). We further analyzed by NGS the maturation of cells edited to express the VRC26-UCA HCDR3. A phylogenetic tree was constructed from the most frequently observed sequences in mice engrafted with B cells expressing this HCDR3 (**Fig. 6B**). We observed that these affinity-matured UCA HCDR3 acquired multiple mutations also found in the VRC26 variants isolated from the original donor, CAP256 (Doria-Rose et al., 2014; Johnson et al., 2018). As in that donor, most active hypermutations focused on residues near a disulfide bond found in more mature VRC26-lineage members. The five most frequently observed HCDR3s were characterized for their ability to neutralize CRF250 pseudoviruses (**Fig. 6C-D**). Four of these five improved the potency of the original VRC26-UCA HCDR3, indicating affinity maturation of this UCA. Thus, SOSIP-TM-expressing mRNA-LNP-delivered immunogens can elicit measurable immune responses in mice engrafted with cells expressing the HCDR3 of multiple apex bnAbs. In doing so, they enable quantitative comparison of candidate HIV-1 vaccines and help to define HCDR3 motifs likely to bind these immunogens.

**Figure 6.**
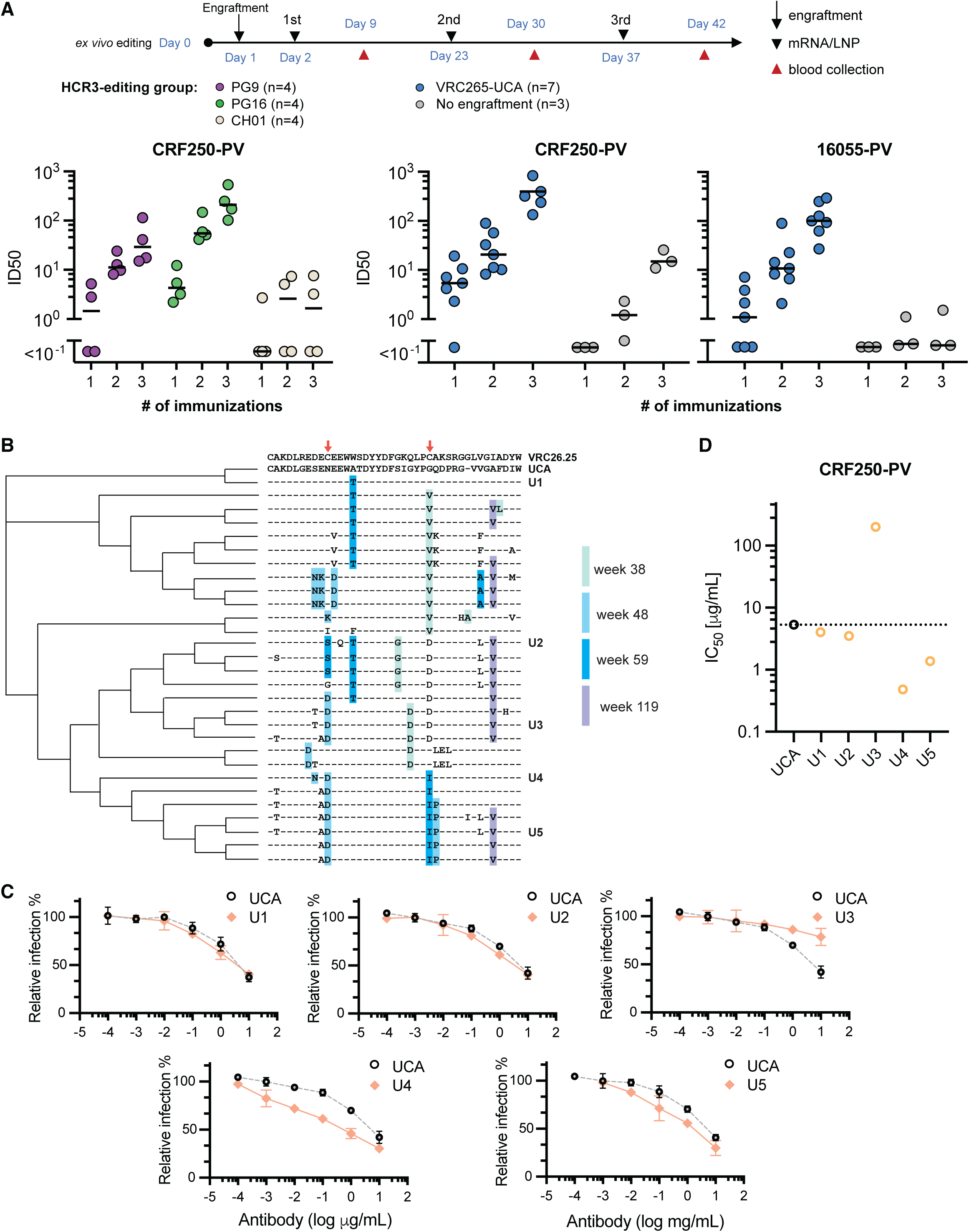
mRNA-LNP elicit neutralizing sera in mice engrafted with B-cells engineered to express the HCDR3s of PG9, PG16, or the VRC26-UCA. **(A)** The indicated numbers of mice per group were engrafted with B cells expressing the HCDR3 of the apex bnAbs PG9, PG16, CH01, and the VRC26-UCA. Serum was harvested one week after each of three immunizations with mRNA-LNP encoding the 16055-ConM-v8.1 SOSIP-TM (PG9, PG16, CH01) or CRF250-ConM-v8.1ds (VRC26-UCA). ID_50_ values were determined using the CRF250 or 16055 pseudoviruses with sera collected from each mouse after the indicated immunization. Grey indicates sera from unengrafted mice immunized in the same manner. **(B)** A phylogenetic tree was constructed from the most frequently observed sequences in mice engrafted with B cells expressing the VRC26-UCA HCDR3. The mature VRC26.25 HCDR3 is shown for reference. The UCA lacks an internal disulfide bond present in the mature HCDR3 (red arrows). The same mutations that also emerged in the human VCR26-lineage were highlighted with different shades of blue, indicating the number of weeks post HIV-1 infection when these mutations first identified. Note the high hypermutation frequency occurred in the vicinity of this missing disulfide bond. The five most frequently observed HCDR3 are indicated with a unique designation (U1-U5). **(C)** The HCDR3 of mature VRC26.25 was replaced by that of VRC26-UCA or the indicated UCA variants and characterized for their ability to neutralize CRF250 pseudoviruses. **(D)** A summary of IC_50_ values calculated from three replicates determined as in panel C is shown.

### Toward an HCDR3-focused strategy for HIV-1 vaccine development

Among the HCDR3 motifs (**Fig. S6A**), the tyrosine-sulfated YYDF sequence derived from D3-3 appears especially important because it is the common motif shared by HCDR3s elicited by SOSIP-TM and because these residues directly engage the Env apex at two sites and in two orientations (**Fig. S6B**), both contributing substantially to Env binding (Pejchal et al., 2010; Andrabi et al., 2015). To determine the frequency of potential apex bnAb precursors, we analyzed long HCDR3s with candidate sulfation motifs from 10 previously characterized human donors (Briney et al., 2019). This analysis defines a potential apex precursor as an antibody with HCDR3 that are 24 amino acids or longer, with a tyrosine-sulfation motif that is 7 residues or more from the beginning of the HCDR3 and 10 amino acids or more from its end. A tyrosine-sulfation motif is broadly defined as at least one tyrosine adjacent to one acidic amino acid, without an adjacent basic residue (Farzan et al., 1999; Monigatti et al., 2002; Choe et al., 2003; Choe and Farzan, 2009). We observed a significant enrichment of candidate precursors for HCDR3 deriving from D3-3, D3-9, D3-16, and D3-22 (**Fig. S6C** and **Table S1**), likely due to the longer lengths of these D gene segments and the presence of sulfation motifs in their dominant reading frames. **Fig. S6D** shows the frequency of these diversity chains and of the specific YYDX sequences encoded by them. An average of one in approximately 200 HCDR3 met these length and sulfation criteria. One in 1800 meet these criteria and encode YYDF. These numbers suggest a considerably higher precursor frequency than for VRC01-class bnAbs, estimated to be 1 in 2.4 million (Jardine et al., 2016). The mouse model described here will help iteratively refine this definition of apex bnAb precursor and help identify SOSIP-TM variants that best prime and affinity mature these precursors.

## DISCUSSION

Here we report a new murine model that can facilitate *in vivo* evaluation of trimeric Env antigens and HIV-1 vaccination strategies, especially those designed to elicit apex bnAbs. This model relies on two distinctive properties of apex bnAbs. First, these bnAbs only bind native-like trimers in a closed conformation. This closed structure occludes immunodominant non-neutralizing epitopes and presents conserved neutralizing epitopes as they are presented by functional Envs on the virion. This model thus provides quantitative insight into the *in vivo* stability and immunogenicity of antigens designed to maintain this structure. A second property of several apex bnAbs, highlighted by our data here, is their near complete reliance on their long HCDR3s to bind Env, providing sufficient affinity to elicit a neutralizing response from diverse murine B cells expressing these HCDR3s. The resulting model is adaptable, sensitive, and efficient, as indicated by its ability to affinity mature the HCDR3s of the bnAb VRC26.25 and its UCA.

This system has a number of advantages over other animal models of vaccination. First, in contrast to previous approaches using transgenic mice (Dosenovic et al., 2015; Escolano et al., 2016; Sok et al., 2016; Tian et al., 2016), it can be established or modified in days, greatly accelerating the developmental cycle of SOSIP and SOSIP-TM antigens. Second, this model enables selection among combinations or libraries of HCDR3s that would be difficult using transgenic mice. Third, in contrast to other murine B-cell editing approaches (Hartweger et al., 2019; Moffett et al., 2019; Voss et al., 2019; Nahmad et al., 2020), this approach does not disrupt the underlying regulatory apparatus at the B-cell locus. Rather, our approach directly overwrites the native HCDR3 without displacing the heavy and light-chain genes or introducing exogenous regulatory elements. Perhaps as a consequence, we observe robust somatic hypermutation and clear affinity maturation of the exogenous HCDR3, not reported with other systems (Hartweger et al., 2019; Moffett et al., 2019; Huang et al., 2020; Nahmad et al., 2020). Fourth, in contrast to most B-cell editing strategies, the underlying BCR diversity is maximized. This underlying diversity better emulates a human repertoire, especially in HIV-1-negative persons, and its preservation has proved valuable in other systems, most notably with transgenic mice expressing human-derived variable chains to evaluate antigens designed to elicit CD4-binding site antibodies (Tian et al., 2016). Fifth, in contrast to wild-type animal models like mice, rabbits, or primates, our system presents human HCDR3s. This property is especially critical because the D gene segments that help form these HCDR3s are highly species specific (Lefranc, 2014; Lefranc et al., 2015). For example, no non-human species has a D segment homologous to human D3-3, D3-9, D3-16, or D3-22. As shown in **Fig. S6**, these D segments are relatively long and directly encode sulfation motifs. Thus *in vivo* testing of antigens designed to elicit human HCDR3 would ideally be performed directly in humans or in the system we present here.

Using this system, we developed several insights useful for generating an apex-targeting vaccine. First, we observed that design differences among SOSIP immunogens significantly altered their ability to elicit apex-focused antibody responses. For example, in **Fig. 4**, we compared multiple SOSIP proteins with identical V1V2 and therefore apex regions, but with different stabilizing mutations. We observed that those bearing v8.1 mutations were significantly more immunogenic (**Fig. 4C**). Notably, the ability of these SOSIP variants to bind HCDR3-edited murine B cells did not fully predict their immunogenicity (**Fig. 4A-C**), underscoring the necessity of *in vivo* models that anticipate human immune responses. We also observed that v8.1 SOSIP variants elicited a less pronounced non-neutralizing response in unengrafted mice (**Fig. 4D-E**). Thus, this closed-form trimer may more effectively occlude non-neutralizing decoy epitopes. Second, our data make clear that mRNA-delivered vaccine candidates expressing these SOSIPs with a transmembrane domain were significantly more immunogenic than soluble SOSIP multimers administered as an adjuvanted protein (**Fig. 5**). Notably, SOSIP variants that elicited more potent responses when presented by mi3-conjuated multimers (**Fig. 4C**) also elicited stronger responses when expressed as SOSIP-TM and delivered by mRNA-LNP (**Fig. 5E**), suggesting that stability improvements useful for soluble SOSIP antigens extend to their use in mRNA vaccines. Third, our data show that SOSIP-TM can elicit neutralizing antibodies from B cells edited to express four distinct HCDR3s, those of PG9, PG16, VRC26.25, and the predicted VRC26 progenitor (**Fig. 5A and 6A**). Thus, the problem of developing a useful – if not comprehensive – vaccine might be reduced to the more tractable goal of eliciting long, sulfated HCDR3 that bind the apices of multiple isolates. Fourth, in contrast to the CH01 HCDR3 which did not generate a neutralizing response in engrafted mice, all HCDR3 that elicited such a response share a sulfated D3-3-encoded YYDF motif (**Fig. S6A**). The HCDR3-edited mouse model described here allows us to test the hypothesis that SOSIPs that most effectively drive maturation of D3-3-expressing HCDR3s, or those deriving from related D segments bearing YYD, can raise apex-binding bnAbs in uninfected humans. As importantly, this model allows us to refine the definition of an apex bnAb precursor by testing additional HCDR3 or HCDR3 libraries derived from human repertoires.

Finally, our data show that repeat immunizations with the same native-like SOSIP-TM immunogen can elicit a neutralizing response against circulating Envs from a bnAb UCA, to our knowledge the first such demonstration. These responses were generated with an initial frequency of 0.2 million transferred UCA-expressing precursor B cells (approximately 0.2% of B cells in a mouse), similar to the frequency of long potentially sulfated HCDR3 observed in humans (0.45%), and to the subset of these HCDR3 containing YYD (0.21%) (**Fig. S6D and Table S1**). Together these observations suggest that SOSIP-TM antigens have the potential to engage these candidate apex precursors to generate a neutralizing response in humans.

In summary, we have developed a useful and adaptable system that can accelerate evaluation and development of candidate antigens for an HIV-1 vaccine. We have also shown that mRNA-expressed SOSIP-TM can generate neutralizing antibodies from B cells edited to express four divergent HCDR3, raising the possibility that BCR with similar HCDR3s can be matured to apex bnAbs in uninfected humans.

## LIMITATIONS OF THE STUDY

This study has several limitations. First, the frequency of potential apex bnAb precursors capable of binding a SOSIP-TM antigen in this model may be greater than in humans. We calculate that the proportion of long, sulfated, and D3.3-derived HCDR3 in humans is higher than the frequency of engineered murine B cells at the time engraftment, but it is unclear how many HCDR3 in a human repertoire can bind a SOSIP-TM antigen with the affinity of the VRC26.25 UCA. A more accurate definition of a human apex bnAb precursor is therefore needed. The system described here can help refine that definition and help evaluate SOSIP-TM variants designed to prime and affinity mature the pool of apex precursors. Second, every described apex bnAb has lower breadth than most CD4 binding-site bnAbs. However, an intact apex implies that other key epitopes also remain intact. Although this model cannot directly monitor the antibody response to these other epitopes, it can accelerate development of SOSIP-TM variants designed to forward the maturation of all major antibody classes. Third, Env does not engage mouse CD4, so potentially destabilizing SOSIP interactions with CD4 in humans will not be modeled. This problem can be mitigated with SOSIP-TM because its transmembrane domain keeps this antigen associated with the LNP-transfected cells, thus limiting these CD4 interactions. Finally, there remain many other differences between mice and humans, including their B-cell repertoires, their MHC and T cell responses, and their *in vivo* environments, that could confound interpretation of results from this system. Future studies that engraft HCDR3-edited B cells into other animal models can confirm conclusions drawn from this mouse model.

## RESOURCE AVAILABILITY

### Lead contact

Further information and requests for resources and reagents should be directed to and will be fulfilled if practicable by the lead contact, Michael Farzan.

### Materials availability

Plasmids generated in this study will be made available under Uniform Biological Material Transfer Agreement (UBMTA) from UF Scripps Biomedical Research.

## EXPERIMENTAL MODEL AND SUBJECT DETAILS

### Mice

Nine to 12 week-old CD45.1-positive mice (B6.SJL-*Ptprc^a^ Pepc^b^*/BoyJ, strain 002014) from Jackson Laboratories were used as a source of splenic B cells. Age- and gender-matched CD45.2-positive C57BL/6J strain mice (Jackson Laboratories, strain 000664) mice were used as host mice for B cell transplantation and immunizations.

All mice were housed and cared at the institutional animal facility in UF Scripps (Jupiter, FL), following the Animal Welfare Act and other federal, state, and local policies and regulations. All animal experiments were reviewed and approved by the Institutional Animal Care and Use Committee under the approval protocol number 21-010-01.

## METHOD DETAILS

### Mouse splenic B cell activation and electroporation

Whole spleens from 9-12 week old CD45.1-positive donor mice were pulverized and mechanically crushed on the inner top of 70 μm cell strainers in RPMI 1640 medium (Thermo Fisher Scientific, 61870127) with 2% FBS (Thermo Fisher Scientific, 26140-079). After red blood cell lysis in a NH_4_Cl solution (BD Biosciences, 555899) at room temperature for 3 minutes, B cells were neutralized with Ca^2+^/Mg^2+^ free PBS with 0.5% BSA (Sigma-Aldrich, A1470) and 5 mM EDTA and then isolated using mouse B cell purification kit (Miltenyi Biotec, 130-090-862) and LS columns (Miltenyi Biotec, 130-042-401). Before electroporation, B cells were activated for 36-42 hours in RPMI 1640 medium with 10% FBS, 100 μM Non-Essential Amino Acids (NEAA, Thermo Fisher Scientific, 11140050), 1 mM sodium pyruvate (Thermo Fisher Scientific, 11360070), 10 mM HEPES (Thermo Fisher Scientific, 15630080), 55 μM 2-Mercaptoethanol (Thermo Fisher Scientific, 21985023), 100 units/mL penicillin and 100 μg/mL streptomycin (Thermo Fisher Scientific, 15140163), and either (1) 4 μg/ml anti-mouse CD180 antibody (BD Biosciences, 562191), (2) 50 μg/ml LPS (Sigma-Aldrich, L2880), or (3) 10 μg/ml LPS and 10 ng/ml mouse IL-4 (PeproTech Inc, 214-14).

After activation, B cells were harvested and washed twice with Ca^2+^/Mg^2+^ free PBS at room temperature. For each 100 μl of electroporation reaction using the nucleocuvette vessels (Lonza, V4XP-4024), approximately 5 million cells were suspended in 74 μl of P4 Primary Cell solution (Lonza, VSOP-4096). In parallel, 3.12 μl of PBS, 1.26 μl of 1M NaCl, 1.12 μl of 250 μM Mb2Cas12a (produced in house), and 4.5 μl of 100 μM gRNA (5’-UAAUUUCUACUGUUUGUAGAUCUUGACCCCAGUAGUCCAUAGCA-3’) were mixed and incubated at room temperature for 15 min for the RNP complex formation. These 10 μl of RNP were then incubated with 16 μl of 100 μM single strand DNA donor (see Table S2 for the sequence details) for 3 minutes at room temperature. The above 26 μl mixture was then mixed with the 74 μl of suspended B cells and transferred to the nucleocuvette vessels for electroporation in the Lonza 4D nucleofector under the DI-100 program. For larger scale electroporation in the 1 ml scale of Nucleocuvette Cartridge (Lonza, V4LN-7002), reactions were scaled up 10 fold for cell suspension, RNP, and donors. After electroporation, cells were rested for 10 minutes in nucleocuvette vessels or cartridges and then transferred to preheated activation medium without penicillin-streptomycin or LPS. which were added one hour later.

### Mouse B cell transplantation

Approximately 18 hours after electroporation, B cells were washed with prechilled Ca^2+^/Mg^2+^ free PBS for four (for LPS activated cells) or three times (for anti-mCD180 antibody activated cells) and then suspended in prechilled 5% horse serum (Cytiva, SH3007403HI) solution in PBS with Ca^2+^/Mg^2+^ (Thermo Fisher Scientific, 14040133). After filtration (Falcon, 352235) and counting, the indicated number of cells for each mouse were transplanted in a 100 μl volume via retro-orbital injection under anesthesia with isoflurane. An aliquot of approximately 2 million cells were further cultivated in RPMI 1640 medium with 10% FBS, 100 μM NEAA, 1 mM sodium pyruvate, 55 μM 2-mercaptoethanol, 10 mM HEPES, 100 units/mL penicillin and 100 μg/mL streptomycin, 5 μg/ml LPS, 10 ng/ml mouse IL-4, and 2 μg/ml anti-mouse CD180 antibody for additional 36-48 hours to monitor editing efficiency validation by flow cytometry.

### Protein production, purification, and conjugation

Protein expression plasmids were constructed using dsDNA gBlocks from IDT and NEBuilder® HiFi DNA Assembly Cloning Kit (NEB, E5520S) and then transformed in NEB 5-alpha competent cells (NEB, C2987U). Expi293F (Thermo Fisher Scientific, A14527) and GnTI-Expi293F (Thermo Fisher Scientific, A39240) cells were maintained in Expi293™ Expression Medium (Thermo Fisher Scientific, A1435102) following the manufacturer’s instructions. Cells were diluted to three million/ml in fresh and preheated medium (Thermo Fisher Scientific, A1435102), and then transfected with FectoPRO reagent (Polyplus, 116-040). To produce SOSIP proteins, plasmids expressing the indicated SOSIP protein, furin, and PDI (protein disulfide-isomerase) were co-transfected at 4:1:1 ratio. For antibody production, plasmids expressing the antibody heavy and light chains were co-transfected at 1:1.25 ratio. In the case of the tyrosine-sulfated PGT145 and VRC26-family antibodies, heavy chain, light chain, and TPST2-expressing plasmids were co-transfected at the ratio for 1.78:2.2:1. For Spc3-mi3-ctag production, Spc3-mi3-Ctag- and PDI-expressing plasmids were co-transfected at the ratio of 4:1. Four to 5 days post-transfection, cell supernatants were harvested, centrifuged, and filtered before purification.

PGT145 and VRC26-family antibodies were captured with Protein A columns (Cytiva, 11003493) and gently eluted with 3 M MgCl_2_ solution (Thermo Fisher Scientific, 21027). Elutions were then buffer exchanged to HEPES buffer (10 mM HEPES pH 8.0 and 75 mM NaCl) firstly and exchanged to PBS with desalting columns (Thermo Fisher Scientific, 89894) following the manufacturer’s instructions. Antibodies were finally concentrated by ultrafiltration. SOSIPs were purified with PGT145 affinity columns and gently eluted with 3 M MgCl_2_ solution, after buffer exchange, trimers were purified by SEC (size exclusion) in the Superdex 200 Increase 10/300 GL column (Cytiva, 28990944) or HiPrep 26/60 Sephacryl S400 HR column (Cytiva, 28935605). Spc3-mi3-Ctag 60-mer were purified with anti-Ctag affinity columns followed with gentle elusion and buffer exchange then the 60-mer were purified by SEC in HiPrep 26/60 Sephacryl S400 HR column. SOSIPs were conjugated onto keyhole limpet hemocyanin carrier protein with Imject EDC mcKLH Spin Kit (Thermo Fisher Scientific, 77671) in accordance with the manufacture’s protocol.

For conjugation to the mi3 60-mer using the SpyTag/SpyCatcher system, purified SOSIP trimers with C-terminal Spy-tag-2 (ST2) and Spc3-mi3-Ctag 60-mer with N-terminal SpyCacher-3 (Spc3) were conjugated at molar ratio of 2:1. Conjugated SOSIP-mi3 multimers were then purified from the free SOSIP-trimers by SEC in HiPrep 26/60 Sephacryl S400 HR column. Conjugation fractions from SEC were pooled, concentrated, and validated with electrophoresis in native, reducing, and non-reducing denaturing SDS-PAGE gels.

### mRNA lipid nanoparticle production

Codon-optimized genes encoding SOSIP variants fused to the Env C-terminal transmembrane domain (TM) sequence were inserted into a pUC vector with 5’ UTR, 3’ UTR, and polyA sequences under T7 promotor. For *in vitro* transcription (IVT), the DNA templates were linearized by digestion with HindIII and ScaI (NEB) and purified by phenol-chloroform extraction. IVT was then performed using MEGAscript® T7 Transcription Kit (Thermo Fisher Scientific, AMB-1334-5) according to the manufacturer’s instructions with modifications as using the CleanCap® Reagent AG (TriLink, N-7413) and m1-pseudouridine-5’-triphosphate (TriLink, N-1081). Template DNA was digested with Turbo DNase, and synthesized mRNA was purified by LiCl precipitation and 75% ethanol washing. After RNA qualification via electrophoresis in a denaturing agarose gel, double stranded RNA was then removed by cellulose (Sigma-Aldrich, C6288) depletion. The mRNA solution was then precipitated with 3M sodium acetate pH 5.2 and washed with isopropanol and then 75% ethanol. Finally, the RNase free water suspended mRNA were quantified and stored at −80°C before LNP formulation.

mRNA LNP were formulated via mixing cartridges in the NanoAssembr BenchTop instrument (Precision, NIT0055) according to the manufacturer’s instructions. First, mRNA was diluted to 0.1-0.35 mg/ml in RNase free water with 25 mM sodium acetate pH 5.0 as the aqueous phase. Corresponding amount of lipid phase, which were one third in volume as the aqueous phase, were calculated with N:P ratio of 6:1 and prepared by adding the lipid solutions SM-102 (MedChemExpress, HY-134541), DSPC (Avanti, 850365), cholesterol (Sigma-Aldrich, C8667), and PEG2000 PE (Avanti, 880150) at the molar ratio of 50:10:38.5:1.5 into ethanol. Aqueous phase and lipid phase were then transferred into individual syringes and loaded to the pre-washed NanoAssemblr Benchtop Acetone Cartridge (Precision, NIT0058). LNP were formulated by mixing of the aqueous phase and lipid phase at a flow ratio of 3:1 and a flow speed of 6 ml/min. After formulation, LNP were buffer exchanged to PBS by dialysis and concentrated via ultrafiltration. mRNA encapsulation efficiencies and concentrations were determined with the Quant-iT RiboGreen RNA Assay Kit (Thermo Fisher Scientific, R11490). Diameters of LNP were measured by dynamic light scattering (DLS) using a Dynapro Naostar (Wyatt Technologies) and, finally, LNP were sterilized by filtration and stored at −80°C in PBS with 10% sucrose.

### Immunizations, blood collection, and B cell isolation for recipient mice

Immunizations were initiated at 24-48h after adoptive transfer of B cells. For protein immunization, indicated dose of conjugated or free SOSIP proteins (25 μg mi3-conjugated SOSIP per mouse in most experiments) were mixed with 20 μg MPLA (Invivogen, vac-mpls) and 10 μg Quil-A (Invivogen, vac-quil) in PBS for a total volume of 250 μl for each mouse. This antigen mixture was injected subcutaneously (s.c.) at four sites (two 50 μl injections underneath two inguinal skin, one 50 μl injection underneath abdomen skin, and one 100 μl injection underneath upper back skin). For mRNA LNP immunizations, the indicated dose of mRNA LNP (5 μg per mouse for most experiments) were diluted in 40 μl volume for each mouse, 20 μl injected into gastrocnemius muscle of each leg. Boost immunizations were administered 2 or 3 weeks, as indicated, for each experiment. Blood samples were collected one week after each immunization via submandibular bleeding. Four days after the final boost, mice were sacrificed and B cell were isolated from spleen and lymph nodes with mouse Pan B Cell Isolation Kit II (Miltenyi Biotec, 130-104-443) and LS columns (Miltenyi Biotec, 130-042-401).

### Pseudovirus production and neutralization assays

HIV envelope plasmids were transformed and amplified in NEB-stable competent cells (NEB, C3040H). Pseudoviruses were produced by co-transfection of plasmids encoding various HIV-1 Envs together with NL4-3 ΔEnv or Q23-ΔEnv (1:3 ratio) in HEK293T cells using PEIpro (Polyplus, 101000033). Plasmids were acquired through the NIH HIV Reagent Program. Supernatant was harvested 48h post transfection, clarified by centrifugation and filtration with a 0.45 µm filter, and aliquoted for storage at −80°C. TZM-bl neutralization assays were performed as previously described. Briefly, titrated mouse sera or antibodies in 96-well plates were incubated with pseudotyped viruses at 37°C for 1 hour. TZM-bl cells were then added to the wells at 10,000 cells/well. Cells were then incubated for 48 hours at 37°C. At 48h post infection, cells were lysed in wells and subjected to firefly luciferase assays. Luciferase expression was determined using the Britelite Plus (PerkinElmer, 6066761) substrate and measured with a Victor Nivo plate reader (PerkinElmer).

### ELISA

To monitor the SOSIP-binding activity of antibodies in mouse sera by indirect ELISA, 96-well plates (Corning, 3690) were coated overnight at 4°C with purified 16055ConM-v8.1ds SOSIP trimers at a concentration of 5 μg/mL in PBS. Wells were washed three times with 0.05% Tween 20 in PBS, and blocked for 1 hour at room temperature with 100 μL of 3% globulin free BSA (Sigma-Aldrich, A7030). After blocking, wells were loaded with 50 μl of serially diluted mouse sera in triplicates for one hour at RT followed by five washes. Wells were then incubated with 50 μL of Fab-specific, peroxidase-labeled goat anti-Mouse IgG (Sigma-Aldrich, A9917) with the dilution for 1:2000 in 1% BSA PBS solution for 30 minutes at RT. Following incubation and five washes, 50 μL of Ultra TMB-ELISA substrate (Thermo Scientific, 34028) was added to each well and incubated at RT for 3 minutes and the reaction was terminated with 50 μL of TMB Stop solution (SouthernBiotech, 0412-01). Absorbance at 450 nm (OD450) was measured with a Victor Nivo plate reader.

For competition ELISA experiments, 96-well plates (Corning, 3690) were coated overnight at 4°C with purified 16055-ConM-v8.1ds SOSIP trimer at a concentration of 5 μg/mL in PBS. Wells were then washed three times with 0.05% Tween 20 in PBS and blocked for 1 hour at room temperature with 100 μL of 3% globulin free BSA (Sigma-Aldrich, A7030) in PBS. Mouse sera was diluted 30-fold with 1% BSA in PBS and 50 μL of diluted sera was added in triplicate to the BSA-blocked wells. Following one-hour incubation with the mouse sera, 50 μL of 1 μg/mL of VRC26.25 or germline-VRC26 was added and mixed for the competitional binding to SOSIP trimers. Incubation of 30 minutes at RT was followed by five washes with 0.05% Tween 20 in PBS. Wells were then incubated with 50 μL of Fc specific, peroxidase labeled goat anti-human IgG (Sigma-Aldrich, A0170) with the dilution of 1:2000 in 1% BSA solution for 30 minutes at RT, and analyzed as described for indirect ELISA studies.

### Staining, sorting, sequencing, and analysis of the mouse B cell IgH repertoire

Engineered mouse B cells were analyzed by double-staining with FITC labeled anti-mouse IgM antibody (Miltenyi Biotec, 130-095-906) and a SOSIP protein labeled with APC using a fluorochrome using the Lightning-Link (R) Fluorescein Antibody Labeling Kit (Novus, 705-0030). For NGS analysis of the gene-editing events, mouse B cells were sorted with SOSIP proteins. For time course studies, mice were sacrificed at the indicated time points to harvest B cells from spleen and lymph nodes. The following reagent and antibody panel was used for flow cytometry analysis: DAPI (Biolegend, 422801), anti-CD19-PerCP/Cyanine5.5 (BioLegend, 152405), anti-CD45.1-FITC (BioLegend, 110706), anti-CD38-APC/Cyanine7 (BioLegend, 102727), anti-GL7-PE (BioLegend, 144607), and APC labeled SOSIP trimers. To analyze the repertoires of edited B cells engineered BCR, antigen-positive mouse B cells were sorted for NGS analysis. Specifically, mouse B cells were first isolated from spleen and lymph nodes using the Pan B-cell Isolation Kit II (Miltenyi Biotec,130-104-443) following the manufacturer’s protocol. Then isolated B cells were labeled with biotinylated anti-CD45.2 antibody (BioLegend, 109803), and then depleted with the anti-biotin Microbeads Ultrapure (Miltenyi Biotec, 130-105-637). Enriched B cells were finally sorted by DAPI, anti-CD45.1 antibody, and SOSIP proteins.

Sorted B cells were lysed for RNA extraction by the RNeasy Micro Kit (Qiagen, 74004). Primers used for reverse transcription and library amplification are provided in Table S2. The IgH repertoire library was prepared as previously described. Briefly, first-strand cDNA synthesis was performed on 11 μl of total RNA using 10 pmol of each primer in a 20 μl total reaction with SuperScript III Reverse Transcriptase (Thermo Fisher, 18080093) according to the manufacturer’s protocol. DNA synthesis reaction was carried out in 100 μl using 10 pmol of each primer with HotStarTaq Plus DNA Polymerase (Qiagen, 203603). Purified dsDNA products were amplified with 10 pmol of each primer in a 100 μl total reaction volume, again with HotStarTaq Plus polymerase. Final indexing was prepared using the NEBNext Multiplex Oligos for Illumina (NEB, E7710S). All PCR products were purified by ExoSAP-IT reagent (Thermo Fisher, 78205.10.ML) and SPRI beads (Beckman Coulter Genomics, SPRIselect). Bead-purified libraries were sequenced using an Illumina MiSeq 2×250 bp paired end reads. Sequencing reads were processed and analyzed as follows: Paired-end reads were quality filtered and trimmed by Trimmomatic, then merged with PANDAseq using the default algorithm. Merged reads were collapsed by UMI through Migec using the “checkout” algorithm. Processed reads were mapped and annotated by Abstar or Mixcr.

### Analysis Software

Analyzed data and sequences were plotted and graphed using python and GraphPad Prism 9. Flow cytometry data were processed using FlowJo 10. GraphPad Prism 9 was used for data analysis.

### Quantification and Statistical analysis

Statistical information including n, mean, median and statistical significance values are indicated in the text or the figure legends. GraphPad Prism 9.01 was used to calculate significant difference by one- and two-way ANOVA, and by paired t test. Data were considered statistically significant at *p < 0.05, **p < 0.01, ***p < 0.001, and ****p<0.0001.

For all pseudovirus neutralization assays, the IC50 (the concentration of mAb needed to obtain 50% neutralization against a given pseudovirus) was calculated from the non-linear regression of the neutralization curve.

## ACKNOWLEDGEMENTS

This work was supported by National Institutes of Health awards U19 AI149646 (M.F.); R21 AI152836 (M.F.); and R01 DA056771 (M.F.).

## COMPETING INTERESTS

W.H., T.O., Y.Y., and M.F. are inventors on a pending patent application describing methods for editing B cells. The authors have no other competing interests.

## AUTHOR CONTRIBUTIONS

W.H., T.O., and M.F. designed this study. W.H., T.O., N.S., N.B., Y.G., X.Z., A.P and X.L. performed experiments. T.O., C.C.B, V.K., A.E.A, and D.M provided bioinformatic analyses. *In vivo* studies were performed by W.H., T.O, N.S., and X.Z., Y.Y., D.M., B.Q, C.C.B., R.W.S., and H.C. provided critical insight, resources, and analysis. W.H., T.O., and M.F. directed experiments, analyzed data, and wrote the manuscript.

## SUPPLEMENTARY FIGURES

**Figure S1. Optimization of HCDR3 editing.** (**A**) The HA tag-encoding homology-directed repair template (HDRT) used in Figure 1C was modified with one to three phosphorothioate (PS) bonds at the 5’ or 3’ end or both, as indicated. These HDRT were used to introduce the HA tag into the HCDR3 region of primary murine B cells. Cells were then analyzed by flow cytometry with an anti-HA (vertical axis) and anti-mouse IgM antibody (horizontal axis). The percentage of HA-positive cells (dark red) is indicated in each panel. (**B**) A summary of three independent experiments similar to that shown in panel A is shown. Arrow indicates the two 3’ PS modification used in all other figures. (**C**) Cells isolated from mouse spleens were activated for 36 to 42 hours with anti-CD180 (4 µg/ml), high dose LPS (50 µg/ml), or low dose LPS (10 µg/ml) and IL-4 (10 µg/ml). Cells were then electroporated with Mb2Cas12a RNP and HDRT encoding an HA-tag or the VRC26.25 HCDR3, replacing endogenous murine HCDR3s, and incubated in the same conditions for another 18 hours. Cells were then analyzed by flow cytometry with an anti-mouse IgM antibody (horizontal axis) and either an APC-labeled anti-HA tag antibody or an APC-labeled SOSIP trimer. Cells activated with anti-CD180 and electroporated with control templates served as a staining control. Numbers indicate the percentage of successfully edited cells. High dose LPS was used in all other figures except where explicitly stated.

**Figure S2. NGS analysis of donor murine B cells** (**A**) NGS analysis of IgH genes of donor CD45.1 cells isolated as in Fig. 2E, except that the distribution of VH1 family genes is shown for donor cells that expressed the native murine HCDR3. Figure shows results combined from donor B cells harvested from two mice for each time point. Note that an enrichment of VH1-64 was observed, similar to the result of successfully edited donor cells. (**B**) A time course experiment similar to Fig. 2A conducted using the CRF250-v7ds conjugated to KLH as immunogens. Two mice were sacrificed at the indicated time points following each immunization. Bar graph shows the frequency of HCDR3 mutations present in unique sequences successfully edited with the HCDR3 of VRC26.25. The mutations also observed in Fig. 3A are marked with triangles. (**C**) At the top, nine individual mutations identified from Fig. 3A are shown beneath the amino acid sequence of the input VRC26.25 HCDR3. Unmodified VRC26.25 or VRC26.25 variants bearing these mutations were expressed as transmembrane antibodies on 293T cell surface and analyzed by flow cytometry with ConM SOSIP proteins (v8.1ds) modified with the CRF250 or BG505 V1V2 regions, as indicated, conjugated to APC. Cells were co-stained with FITC-labeled anti-human Fc. The ratio between these two signals is shown. (**D**) A figure similar to that in panel **C** except that the indicated combinations of mutations were analyzed. The neutralization profiles of the highest binding variants (NEE, NER, and NERE) are shown in Fig. 3B and **C**.

**Figure S3. *In vitro* down-selection of SOSIP variants** (**A**) Linear representations of the indicated SOSIP versions (v5, v7ds, v8.1ds, v8.1 and v8.1mut3ds). Constant and variable regions of Env gp120 and the helical repeat (HR) regions of Env gp41 are indicated, Mutations shown in blue indicate modifications from v5 SOSIP sequence to v7 SOSIP, red indicates modifications from v7 to v8.1, and green indicated modification from v8.1 to v8.1mut3. An additional ‘ds’ disulfide bond is indicated with the extra bracket linking I201C to A433C. The positions of “V1V2” region and the “base” region were labeled with HXB2 numbering. These SOSIP variants were generated using the base from BG505, CRF504, or ConM HIV-1 isolates, which in many cases were modified by a V1V2 region from a different isolate. Thus, 16055-ConM-v8.1 indicates a ConM-v8.1 SOSIP modified with the V1V2 region of the 16055 isolate. (**B**) Sequence of the V1V2 region of Envs used to modified on different bases. (**C**) The indicated SOSIP variants were characterized for their ability to bind primary murine B cells edited to express the VRC26.25 HCDR3. Each variant is labeled according to the HIV-1 isolate contributing its V1V2 region (BG505, CRF250, ConM), the remainder of the engineered Env ectodomain (BG505, ConM), and the SOSIP version used (v5, v7ds, v8.1, v8.1ds). Protein was produced in Expi293F cells or the same cells lacking the acetylglucosaminyltransferase enzyme (GnTI-). Numbers indicate the percentage of SOSIP-binding cells observed in the indicated gate. (**D**) The percentage of SOSIP-binding cells and mean fluorescent intensity of these cells analyzed in panel **B** are represented.

**Figure S4. Optimization of a murine model engrafted with VRC26.25 HCDR3-edited B cells** (**A**) An SEC-purification profile of Expi293F expressed ConM-v8.1ds SOSIPs conjugated on mi3-60mers using a Sephacryl S-400 HR HiPrep 26/60 column. The elution fractions of the SOSIP-mi3 and unconjugated SOSIP are indicated. (**B**) SDS-PAGE analysis of the indicated conjugated and unconjugated SOSIPs before and after SEC purification, stained by Coomassie blue. The molecular weight of marker is included. ST2 indicates the presence of a SpyTag2 at the SOSIP C-terminus; SPC3 indicates a SpyCatcher3 domain at the N-terminus of the mi3 60-mer. (**C**) 16 µg of the ConMv8.1ds SOSIP trimers conjugated to KLH, the mi3 60-mer, or as a free trimer (none). (**D**) mi3-conjuates of the 16055-ConM-v8.1ds or BG505-ConM-v8.1ds SOSIP variants were produced in Expi239F cells or the same cell lacking the enzyme GnTI (GnTI-) which adds complex glycans on SOSIP proteins. (**E**) Varying amounts of MPLA adjuvant and protein antigens were tested. Two protein antigens (16055-ConM-v8.1ds and BG505-ConM-v8.1ds) were tested in different ranges of dosage. (**F**) Different *ex vivo* activation methods (anti-CD180, high-dose LPS) of B cells and number of donor B cells transferred to the recipient mouse were tested. For panel **C-F,** immunization efficiency was determined by the neutralizing response in mice engrafted with HCDR3-edited B cells. Neutralization assays used sera harvested one week after the second immunization, with the same schedule shown in Fig. 1E. Statistical differences were determined by one-way ANOVA (ns, not significant, *, p<0.05 and **, p<0.01, and ***, p<0.001). (**G**) Mice immunized with 16055-ConM-v8.1ds from Fig. 4H were further characterized. A table summarizing ID_50_ values of pooled sera from neutralization assays against the indicated pseudoviruses derived from tier 2 HIV-1 isolates from the indicated clades. Sera was collected from mice with or without engraftment of edited B cells one week after each immunization. (**H**) Sera from engrafted mice characterized in panel **G** was further harvested each month at the time points indicated, and the ID_50_ of neutralizing sera against the CRF250 pseudovirus was plotted with days.

**Figure S5. Potent immune responses elicited by mRNA-LNP expressing SOSIP-TM variants** (**A**) A linear representation of a SOSIP-TM construct. The membrane proximal region (MPER), the transmembrane domain (TM) and a cytoplasmic domain truncated at residue 712 (CT) are added to the SOSIP constructs shown in **Fig. S3A**. SOSIP-TM nomenclature is identical to that used for soluble SOSIP proteins. (**B**) For the experiment in Fig. 5A, mice were sacrificed after two immunizations to monitor the *in vivo* response of HIV-1 Env-specific donor B cells. Top panel shows flow cytometry analysis of germinal-center B cells (GL7+/CD38-). Bottom panels show the percentage of CD45.1 donor mouse B cells that recognized the HIV-1 Env trimer [SOSIP(+)] within the germinal center. (**C**) The diameter of LNP particles was monitored by dynamic light scattering to quality control individual LNP preparations. The distribution profile shows mRNA-LNP expressing the indicated SOSIP-TM variants. (**D**) The ability of sera from these mice to bind to 16055-ConM-v8.1ds SOSIP protein was determined by ELISA. Colored bars indicate pooled sera from 7 immunized mice engrafted with HCDR3-edited B cells, and gray bars indicated pooled sera from 3 mice immunized without engraftment with edited B cells. Each sample was assayed with three technical replicates. Neutralization studies of the same sera are shown in Fig. 5F. (**E**) Blocking ELISA assays were performed to evaluate the specificity of antibody responses. Specifically, the indicated sera analyzed in panel **D** was used to prevent binding of VRC26.25 or a germline form of VRC26.25 to 16055-ConM SOSIP. The percentage of blocking was calculated based on the binding of VRC26.25 in the presence of naïve mouse serum. Grey indicates vaccinated mice without engraftment with edited B cells. Significant differences in panels **D-E** were determined by two-way ANOVA (ns, not significant; *p<0.05; **p<0.01; ***p<0.001; ****p<0.0001).

**Figure S6. A proposed approach for developing an apex-focused HIV-1 vaccine** (**A**) An alignment of the HCDR3 sequences of PG9, PG16, VRC26.25 and the VRC26-UCA, each deriving from IGHD3-3, and the open reading frames of four D3 diversity segments encoding YYDX. Homology with the YYDF motif present in D3-3 are highlighted in green. Other homologies with D3-3 are indicated in grey. Tyrosines shown to be sulfated are indicate in red. (**B**) Previously reported structures of Env trimers complexed with the bnAbs VRC26.25 (PDB: 6VTT) or PG9 (PDB: 5VJ6, (Wang et al., 2017))are presented with only YYDF region of the HCDR3 of these bnAbs shown. Individual Env protomers are indicated in shades of grey, YYDF contact residues on Env are indicated in light blue. Green, red, blue, and yellow indicate YYDF carbon, oxygen, nitrogen, and sulfur atoms, respectively. (**C**) Previously described BCR repertoires from 10 HIV-1 uninfected human donors were analyzed. The percentage of HCDR3 deriving from the indicated D segment is subtracted from the percentage of the same D segment in the population of ‘apex-like’ HCDR3. A HCDR3 is defined as apex-like if its length is 24 or greater, and it has a tyrosine-sulfation motif 7 or more residues from the HCDR3 amino-terminus and 10 or more residues from its carboxy-terminus. A tyrosine sulfation motif is described as a tyrosine adjacent to an acidic residue without a proximal positive residue. Note that four D3 segments are enriched in the apex-like population. Each of these bears a YYDX motif. (**D**) The frequency of long, sulfated ‘apex-like’ HCDR3 from each donor is shown, along with subsets with bearing the indicated motifs, and subsets deriving from the indicated D3 gene segments.

