## Supplementary material for "Heavy-chain CDR3-engineered B cells facilitate *in vivo* evaluation of HIV-1 vaccine candidates": Fig. S

**Figure S1**

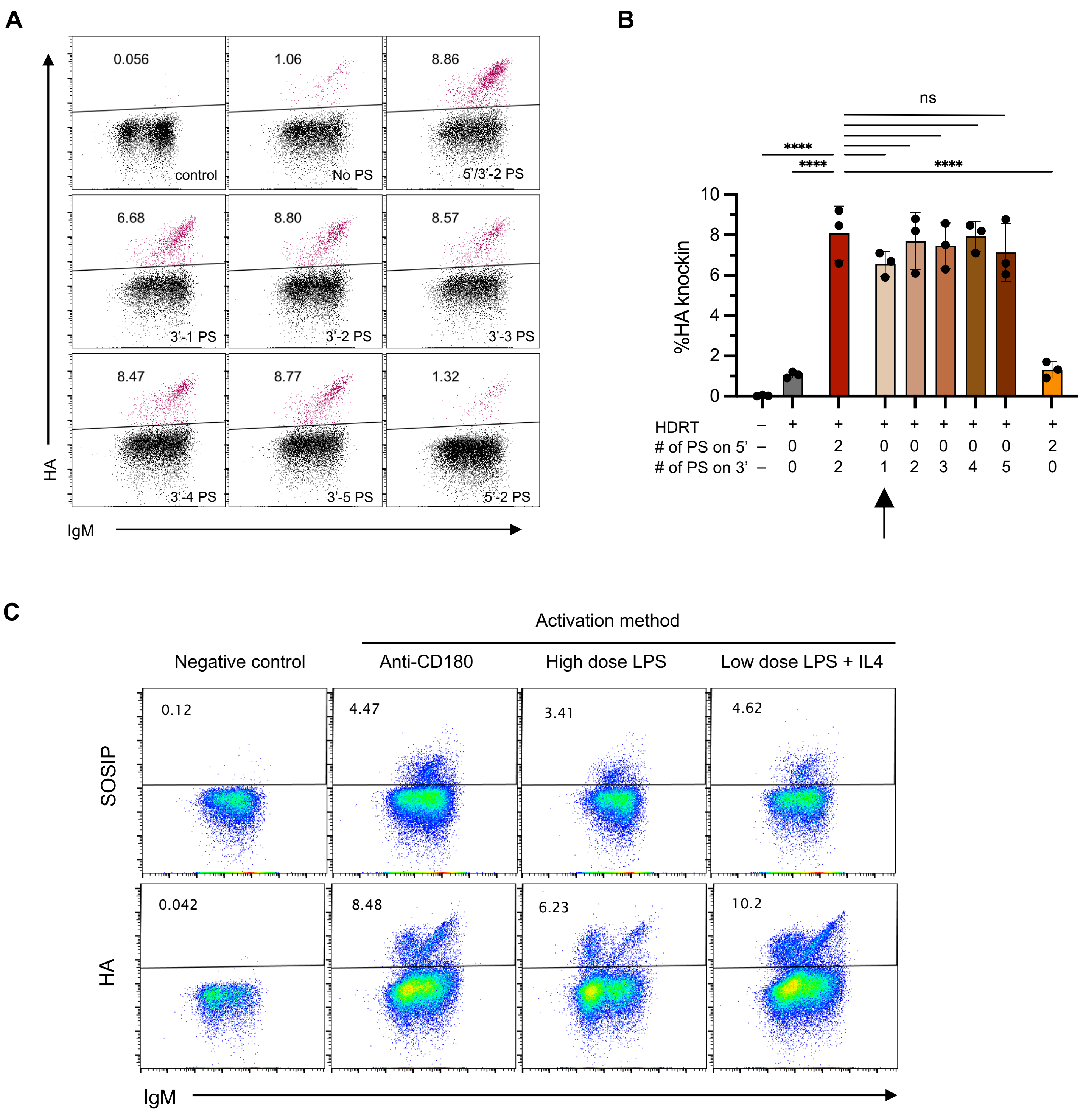
A
0.056
control
1.06
No PS
8.86
5'/3'-2 PS
6.68
3'-1 PS
8.80
3'-2 PS
8.57
3'-3 PS
8.47
3'-4 PS
8.77
3'-5 PS
1.32
5'-2 PS
HA
IgM
B
ns
****
****
****
%HA knockin
HDRT
# of PS on 5'
# of PS on 3'
C
Activation method
Negative control
Anti-CD180
High dose LPS
Low dose LPS + IL4
0.12
4.47
3.41
4.62
0.042
8.48
6.23
10.2
SOSIP
HA
IgM

**Figure S2**

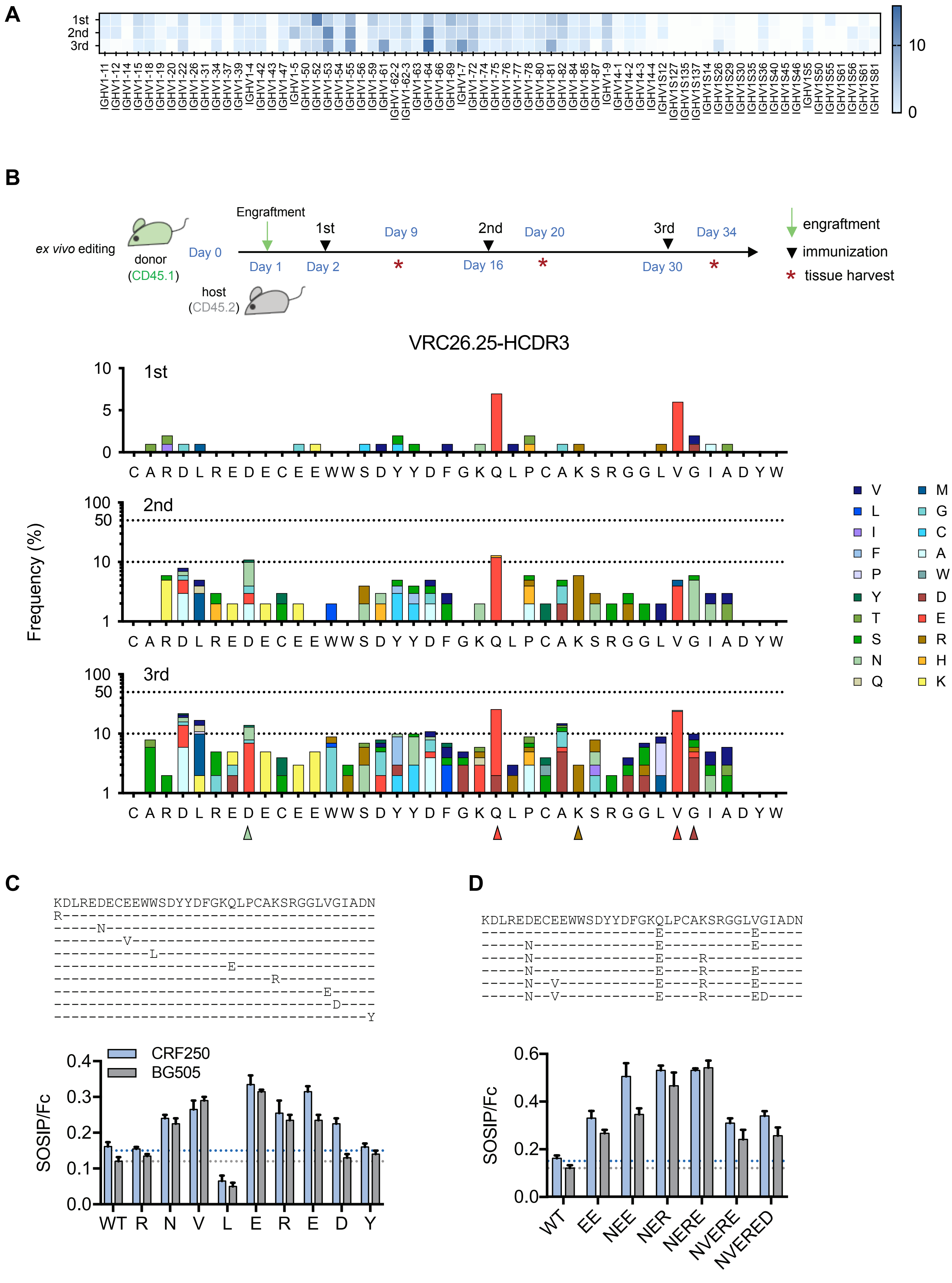
A
1st
2nd
3rd
10
0
IGHV1-11 IGHV1-12 IGHV1-14 IGHV1-15 IGHV1-18 IGHV1-19 IGHV1-20 IGHV1-22 IGHV1-26 IGHV1-31 IGHV1-34 IGHV1-37 IGHV1-39 IGHV1-4 IGHV1-42 IGHV1-43 IGHV1-47 IGHV1-5 IGHV1-50 IGHV1-52 IGHV1-53 IGHV1-54 IGHV1-55 IGHV1-56 IGHV1-59 IGHV1-61 IGHV1-62-2 IGHV1-62-3 IGHV1-63 IGHV1-64 IGHV1-66 IGHV1-69 IGHV1-7 IGHV1-72 IGHV1-74 IGHV1-75 IGHV1-76 IGHV1-77 IGHV1-78 IGHV1-80 IGHV1-81 IGHV1-82 IGHV1-84 IGHV1-85 IGHV1-87 IGHV1-9 IGHV14-1 IGHV14-2 IGHV14-3 IGHV14-4 IGHV1S12 IGHV1S127 IGHV1S135 IGHV1S137 IGHV1S14 IGHV1S26 IGHV1S29 IGHV1S30 IGHV1S35 IGHV1S36 IGHV1S40 IGHV1S45 IGHV1S46 IGHV1S5 IGHV1S50 IGHV1S55 IGHV1S56 IGHV1S61 IGHV1S81
B
ex vivo editing
donor (CD45.1)
host (CD45.2)
Engraftment
Day 0
Day 1
1st
Day 2
Day 9
2nd
Day 16
Day 20
3rd
Day 30
Day 34
engraftment
immunization
tissue harvest
VRC26.25-HCDR3
Frequency (%)
1st
2nd
3rd
C A R D L R E D E C E E W W S D Y Y D F G K Q L P C A K S R G G L V G I A D Y W
V M L G I C F A P W Y D T E S R N H Q K
C
KDLREDECEEWWSDYYDFGKQLPCAKSRGGLVGIADN
R------------------------------------
-----N-------------------------------
---------V---------------------------
------------L------------------------
---------------------E---------------
----------------------------R--------
-----------------------------------E-----
------------------------------------D----
------------------------------------Y
CRF250
BG505
SOSIP/Fc
WT R N V L E R E D Y
D
KDLREDECEEWWSDYYDFGKQLPCAKSRGGLVGIADN
--------------------E----------E-----
-----N--------------E----------E-----
-----N--------------E----R-----------
-----N--------------E----R-----E-----
-----N--V-----------E----R-----E-----
-----N--V-----------E----R-----ED----
SOSIP/Fc
WT EE NEE NER NERE NVERE NVERED

### Figure S3

**A** <u>**SOSIP version:**</u>

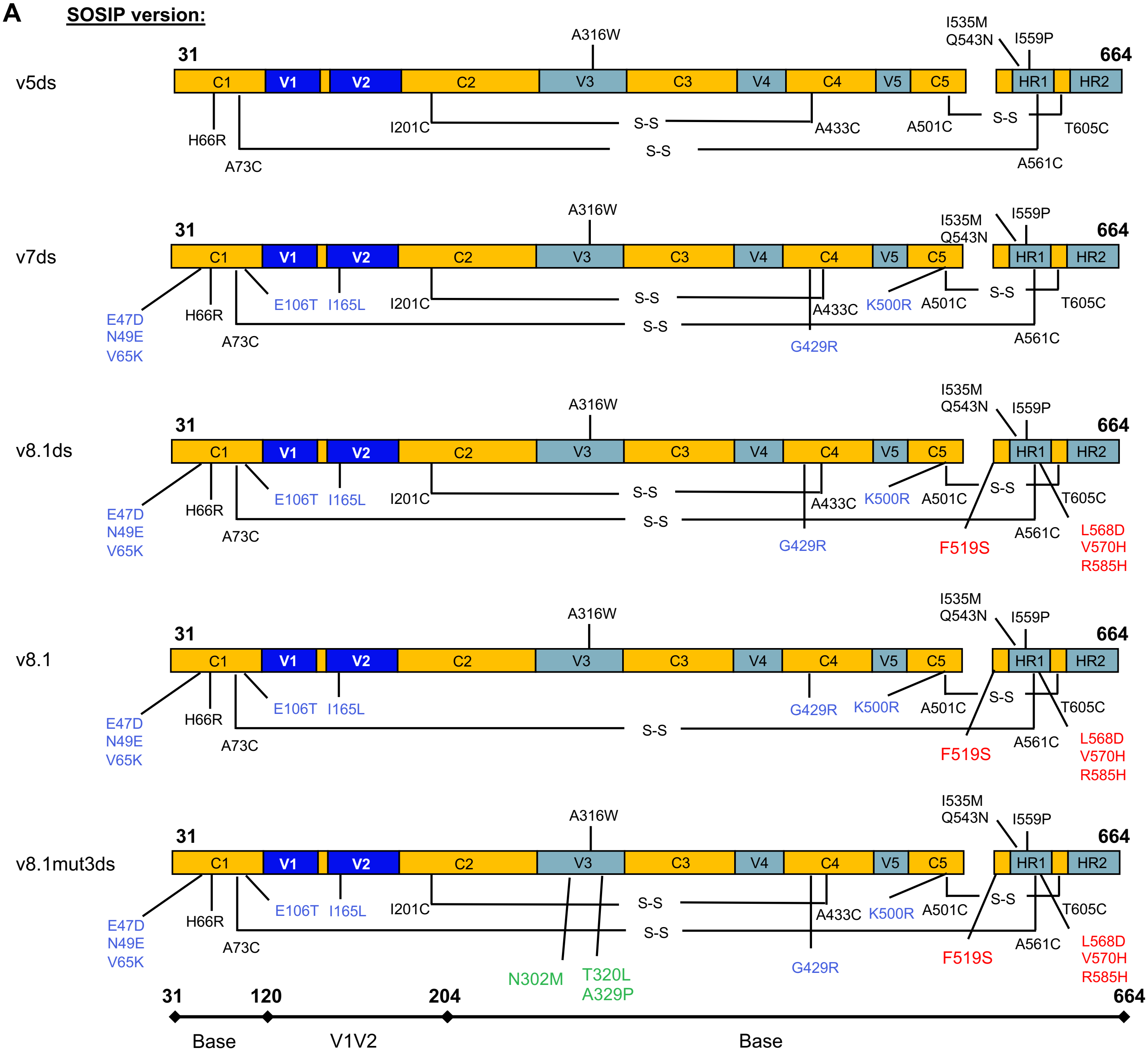


**B** <u>**V1V2 region:**</u>

```
HXB2    120-------130-------140-------150-------160-------170-------180---------------190-------200----
BG505   CVKLTPLCVTLQCTNVTNNITDDM--------RGELKNCSFNMTTELRDKKQKVYSLFYRLDVVQINENQGNRSNNSNKEYRLINCNTSAITQAC
16055   CVKLTPLCVTLECRQVNTTNATS---SVNVTNGEEIKNCSFNATTEIRDKKQKVYALFYRLDIVPLEEERKG----NSSKYRLINCNTSAITQAC
CRF250  CVKLTPLCVTLDCQAFNSSSH-----TNSSIAMQEMKNCSFNVTTELRDKKKKEYSFFYKTDIEQINK--------NGRQYRLINCNTSAITQAC
```

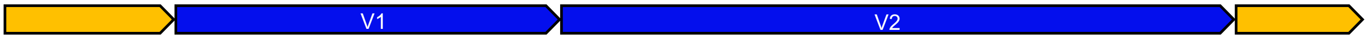

**Figure S3 (continued)**

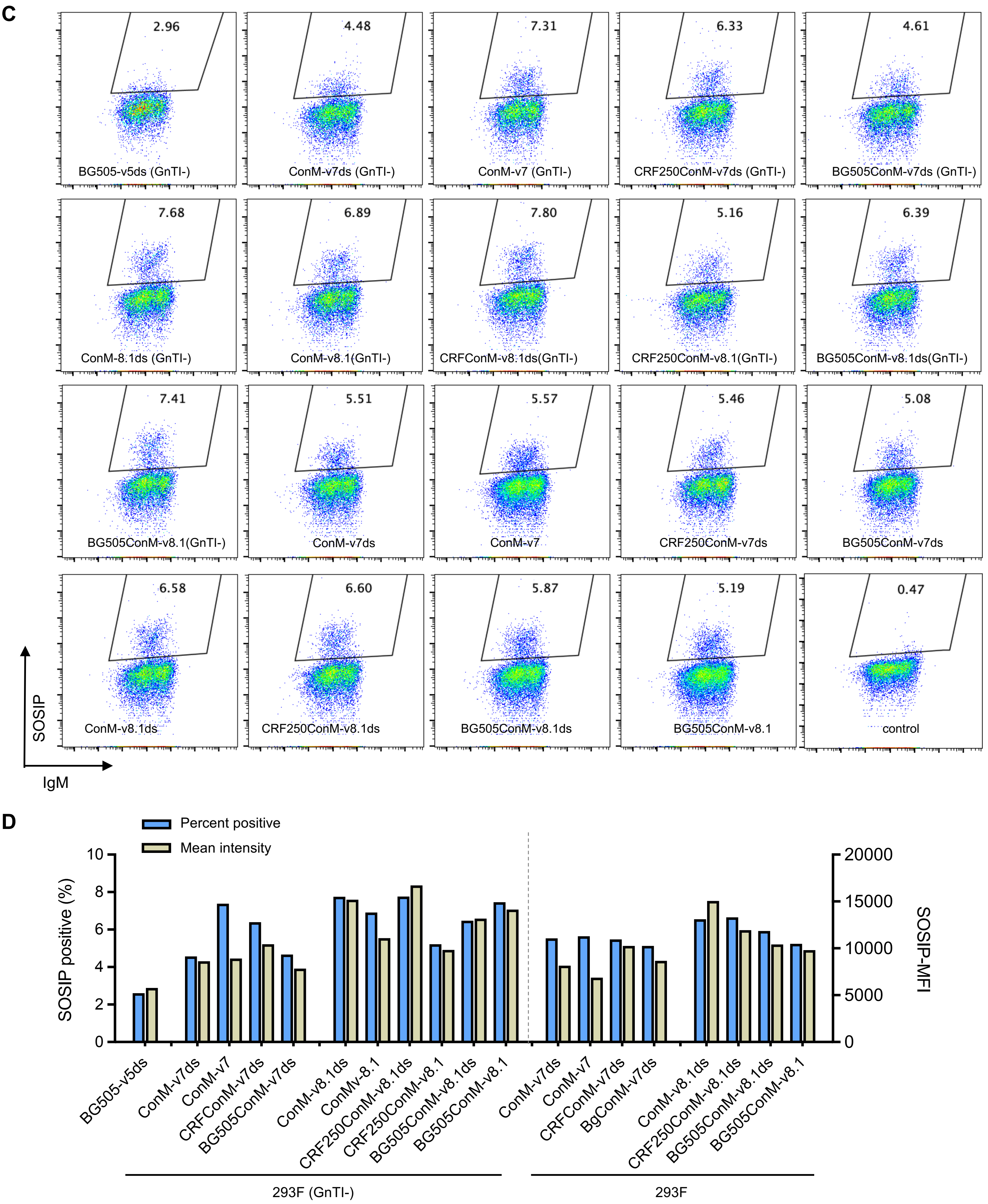

**Figure S4**

**A**

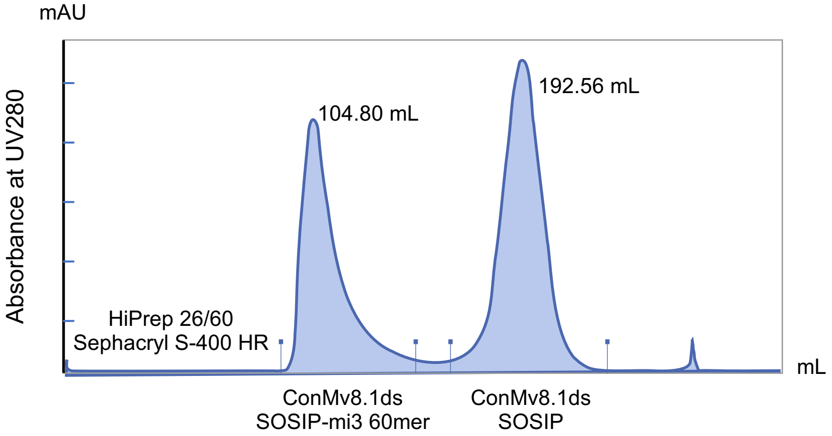


**B**

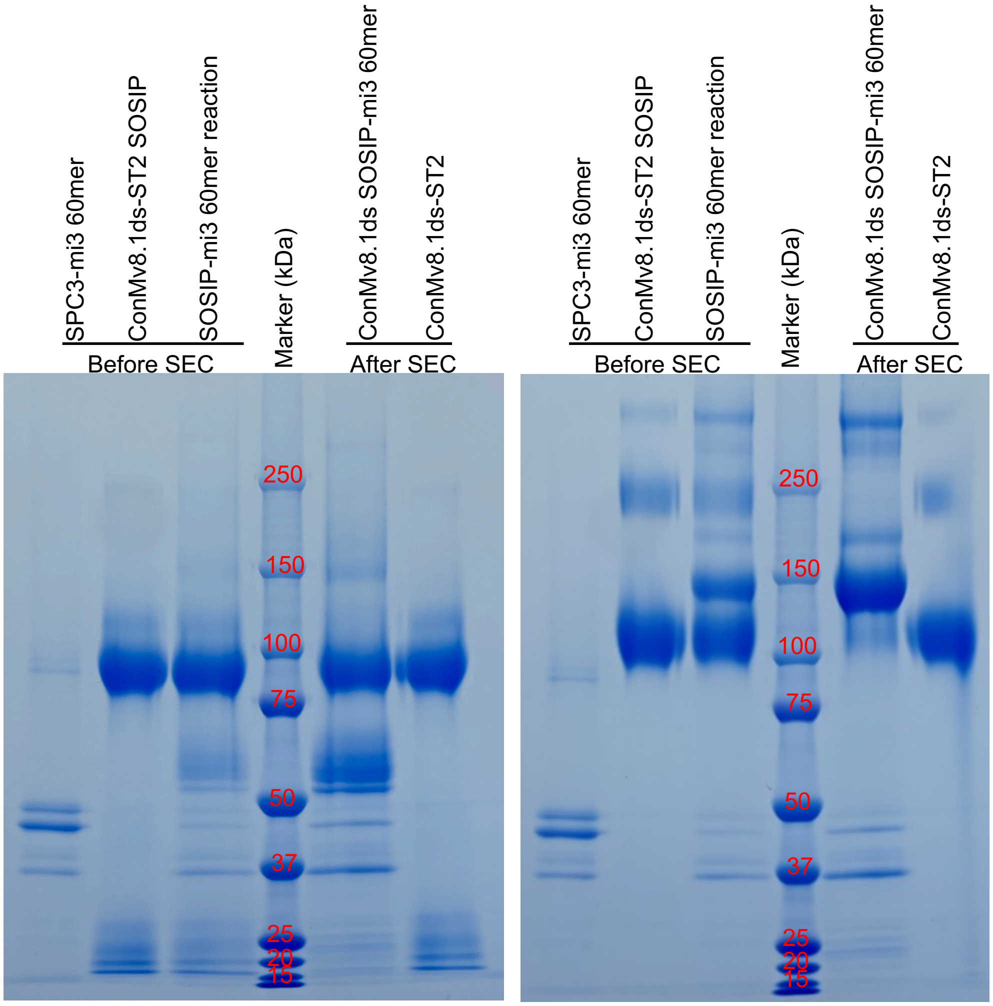


With DTT, reducing SDS-PAGE

Without DTT, non-reducing SDS-PAGE

### Figure S4 (continued)

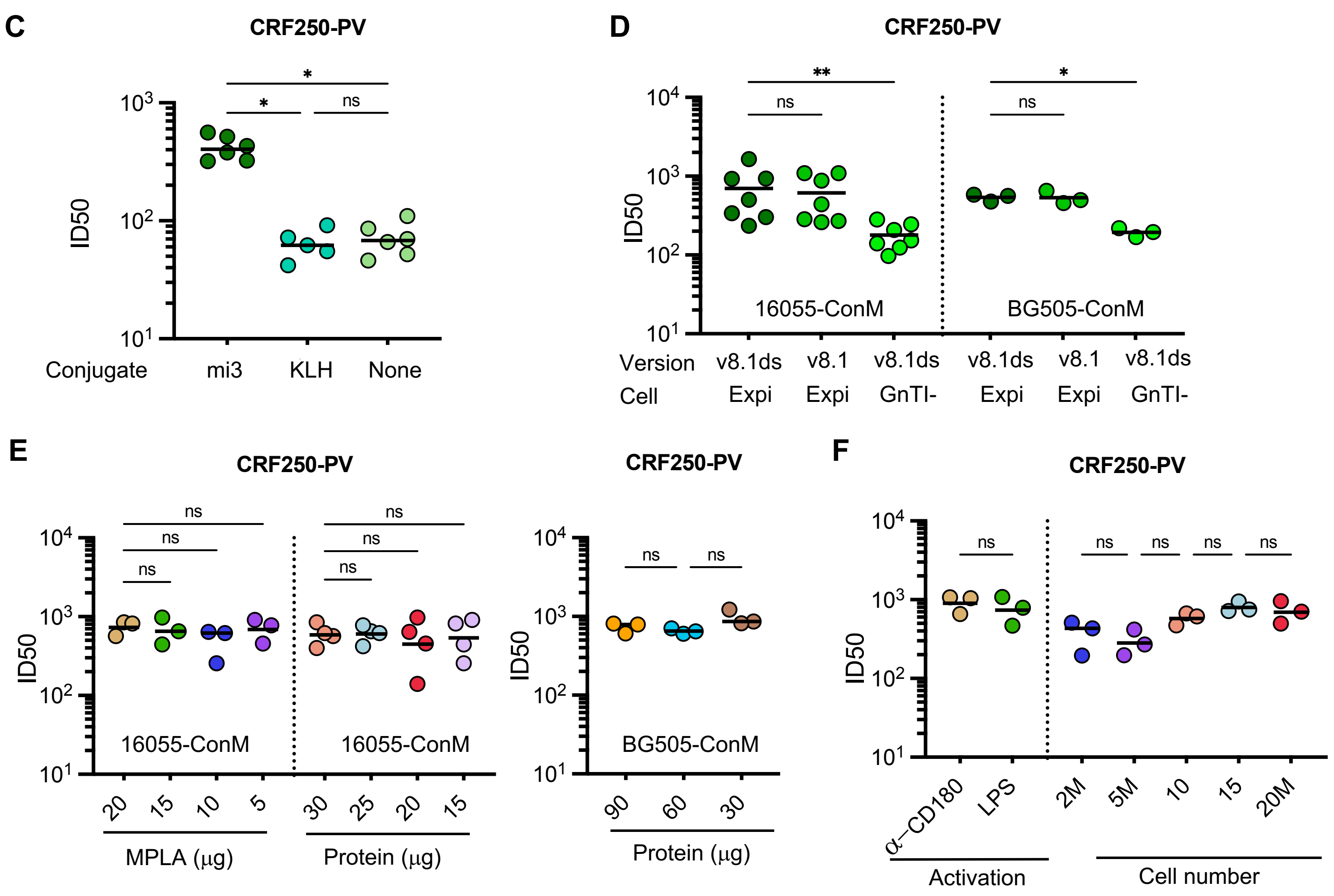


G

| | | 1st immunization | | 2nd immunization | | 3rd immunization | |
|---|---|---|---|---|---|---|---|
| Psudoviruses | Cade | engraft | unengraft | engraft | unengraft | engraft | unengraft |
| BG505 | A | <20 | <20 | <20 | <20 | 23.5 | <20 |
| ZM233 | C | <20 | <20 | 95.2 | <20 | 193.7 | <20 |
| CRF250 | AG | 213 | <20 | 992 | <20 | 4390 | <20 |
| 16055 | C | 102 | <20 | 421 | <20 | 1120 | <20 |
| 25710 | C | 73.2 | <20 | 122 | <20 | 297.5 | <20 |
| x1632 | G | <20 | <20 | <20 | <20 | 26.6 | <20 |
| CNE8 | AE | <20 | <20 | <20 | <20 | <20 | <20 |
| WITO | B | <20 | <20 | <20 | <20 | <20 | <20 |
| % neutralized | | 3/8 | 0/8 | 4/8 | 0/8 | 6/8 | 0/8 |

ID50: >1000; 300-1000; 100-300; 20-100; <20

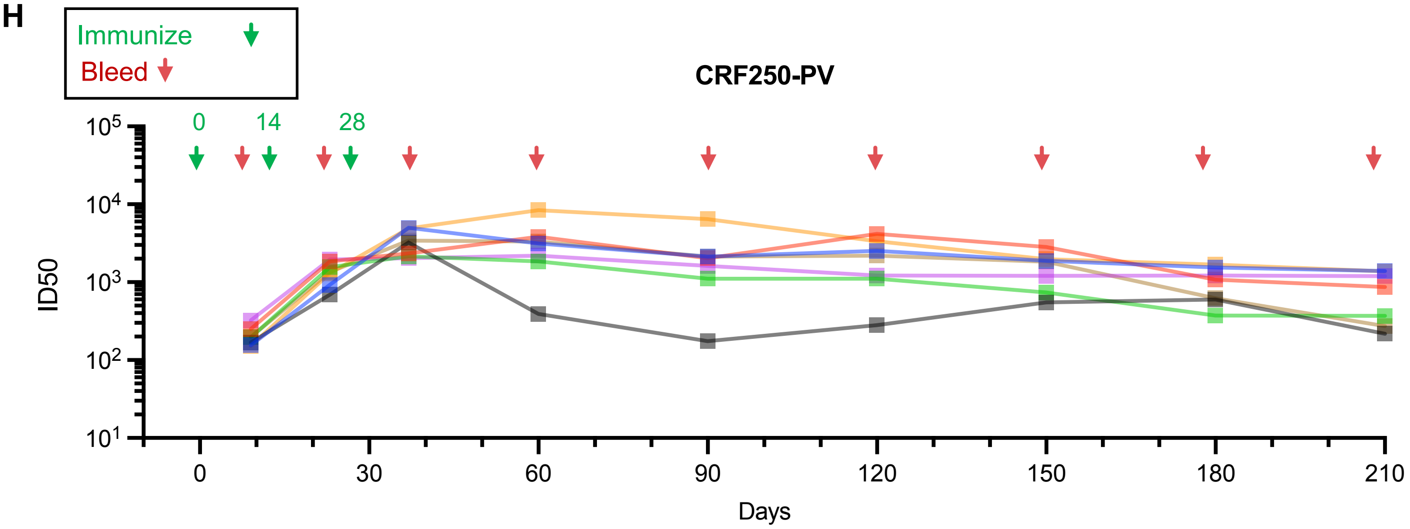

### Figure S5

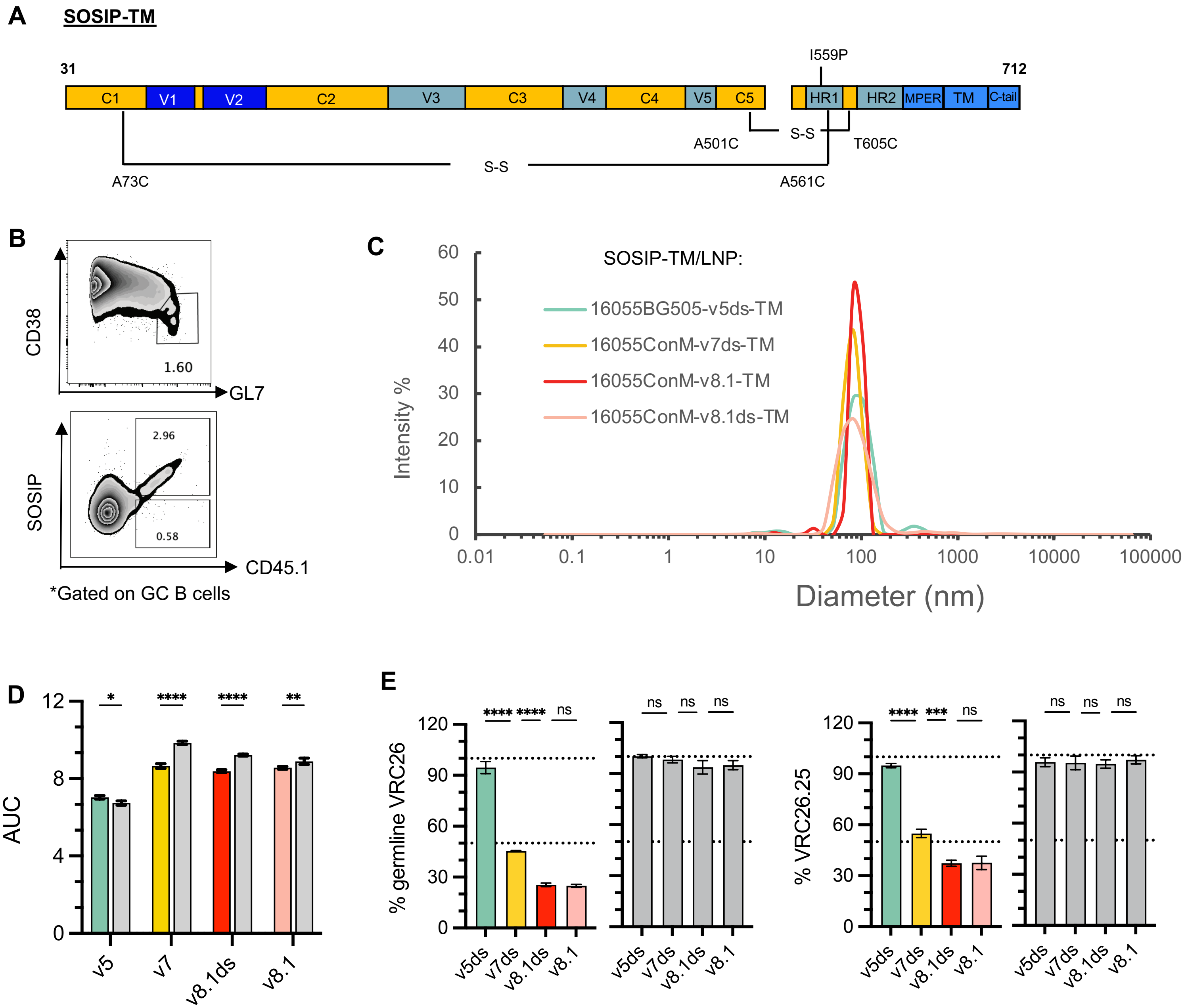

A
SOSIP-TM
31
712
I559P
C1
V1
V2
C2
V3
C3
V4
C4
V5
C5
HR1
HR2
MPER
TM
C-tail
A501C
S-S
T605C
A73C
S-S
A561C
B
CD38
1.60
GL7
SOSIP
2.96
0.58
CD45.1
*Gated on GC B cells
C
SOSIP-TM/LNP:
16055BG505-v5ds-TM
16055ConM-v7ds-TM
16055ConM-v8.1-TM
16055ConM-v8.1ds-TM
Intensity %
Diameter (nm)
D
AUC
v5
v7
v8.1ds
v8.1
E
% germline VRC26
% VRC26.25
v5ds
v7ds
v8.1ds
v8.1

**Figure S6**

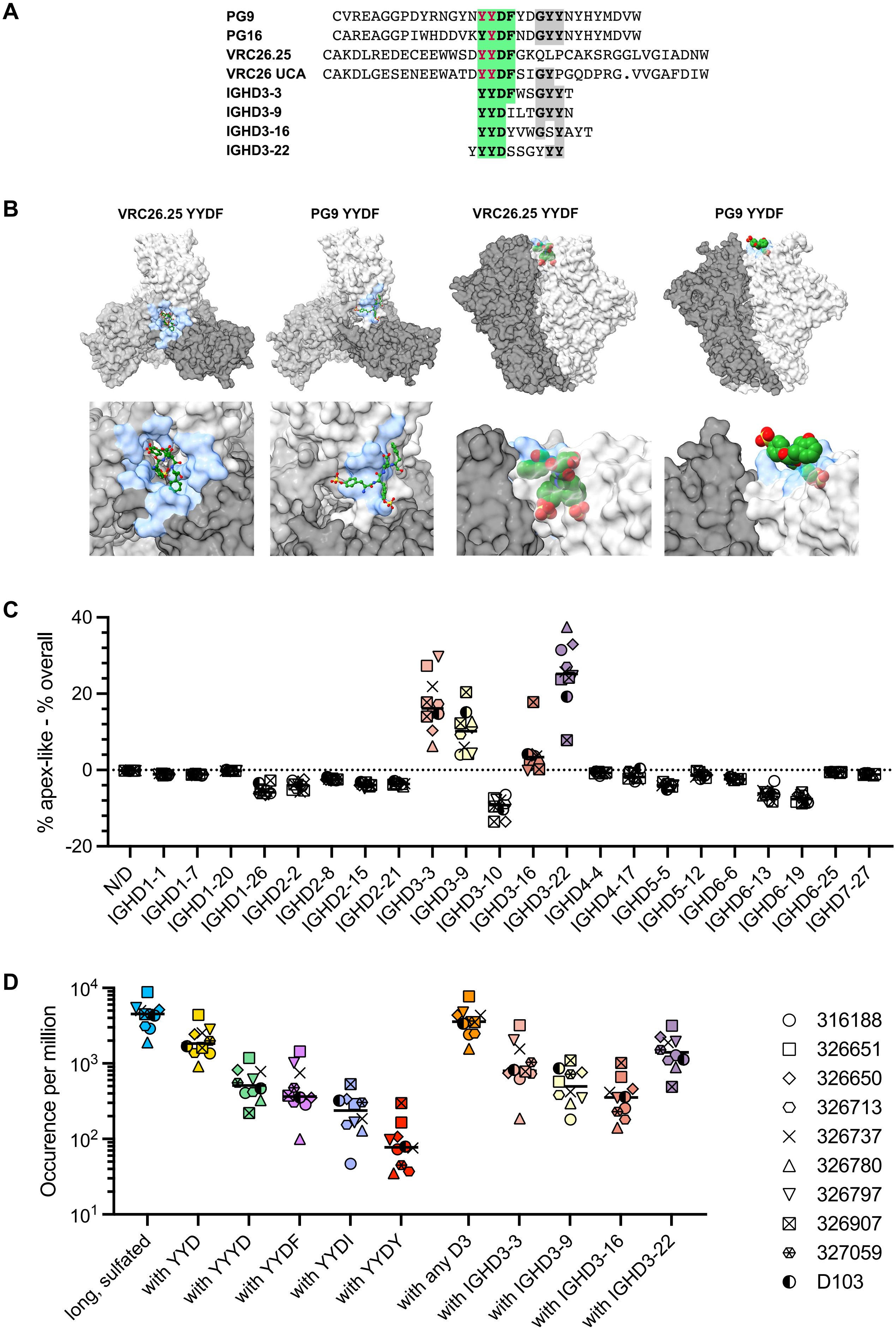

A
PG9 CVREAGGPDYRNGYNYYDFYDGYYNYHYMDVW
PG16 CAREAGGPIWHDDVKYYDFNDGYYNYHYMDVW
VRC26.25 CAKDLREDECEEWWSDYYDFGKQLPCAKSRGGLVGIADNW
VRC26 UCA CAKDLGESENEEWATDYYDFSIGYPGQDPRG.VVGAFDIW
IGHD3-3 YYDFWSGYYT
IGHD3-9 YYDILTGYYN
IGHD3-16 YYDYVWGSYAYT
IGHD3-22 YYYDSSGYYY
B
VRC26.25 YYDF
PG9 YYDF
VRC26.25 YYDF
PG9 YYDF
C
% apex-like - % overall
40
20
0
-20
N/D
IGHD1-1
IGHD1-7
IGHD1-20
IGHD1-26
IGHD2-2
IGHD2-8
IGHD2-15
IGHD2-21
IGHD3-3
IGHD3-9
IGHD3-10
IGHD3-16
IGHD3-22
IGHD4-4
IGHD4-17
IGHD5-5
IGHD5-12
IGHD6-6
IGHD6-13
IGHD6-19
IGHD6-25
IGHD7-27
D
Occurence per million
10^4
10^3
10^2
10^1
long, sulfated
with YYD
with YYYD
with YYDF
with YYDI
with YYDY
with any D3
with IGHD3-3
with IGHD3-9
with IGHD3-16
with IGHD3-22
316188
326651
326650
326713
326737
326780
326797
326907
327059
D103
